# Multi-Channel smFRET study reveals a Compact conformation of EF-G on the Ribosome

**DOI:** 10.1101/2024.01.27.577133

**Authors:** Jordan L. Johnson, Jacob H. Steele, Ran Lin, Victor G. Stepanov, Miriam N. Gavriliuc, Yuhong Wang

## Abstract

While elongation factor G (EF-G) is crucial for ribosome translocation, the role of its GTP hydrolysis remains ambiguous. EF-G’s indispensability is further exemplified by the phosphorylation of human eukaryotic elongation factor 2 (eEF2) at Thr56, which inhibits protein synthesis globally, but its exact mechanism is not clear. In this study, we developed a multi-channel single-molecule FRET (smFRET) microscopy methodology to examine the conformational changes of *E. coli* EF-G induced by mutations that closely aligned with eEF2’s Thr56 residue. We utilized Alexa 488/594 double-labeled EF-G to catalyze the translocation of fMet-Phe-tRNA^Phe^-Cy3 inside Cy5-L27 labeled ribosomes, allowing us to probe both processes within the same complex. Our findings indicate that in the presence of either GTP or GDPCP, wild-type EF-G undergoes a conformational extension upon binding to the ribosome to promote normal translocation. On the other hand, T48E and T48V mutations did not affect GTP/GDP binding or GTP hydrolysis, but impeded Poly(Phe) synthesis and caused EF-G to adopt a unique compact conformation, which wasn’t observed when the mutants interact solely with the sarcin/ricin loop. This study provides new insights into EF-G’s adaptability and sheds light on the modification mechanism of human eEF2.

## INTRODUCTION

The ribosome is a large ribonucleoprotein complex comprising two distinct subunits: a large subunit (LSU) and a small subunit (SSU). It assembles the polypeptide chain, using mRNAs as templates. During protein synthesis, the ribosome moves accurately on the mRNA from 5’ to 3’-terminal, advancing by 3 nucleotides (nt) per step. Within the ribosome, two codons are exposed at positions termed A-site and P-site. The aminoacyl tRNA delivers a new amino acid to the A-site codon, while peptidyl tRNA at the P-site holds the growing peptidyl chain. Following the peptidyl transfer reaction, the peptidyl chain is transferred to the A-site tRNA and lengthened by one amino acid. The tRNAs form codon-anticodon interactions with the mRNA at anti-codon ends and hold the aminoacyl or peptidyl group at the acceptor end, oriented orthogonally, thereby preventing entanglement of the mRNA and peptide chain (1). In this pre-translocation complex (pre-Rb), the peptidyl-tRNA resides at the A-site, while the deacyl-tRNA is situated at the P-site. In the following translocation phase, the tRNAs from the A-/P-sites shift to the P-/E-sites, thereby forming the post-translocation complex (post-Rb). A new vacant codon enters the A-site, and the cycle repeats until a stop codon prompts the termination of the mRNA reading frame. Throughout this process, the mRNA and tRNAs establish extensive interactions with the ribosome (2). Thus, the translocation step involves a series of ribosomal motions to break and form intermolecular bonds (3), such as the 30S head swivel (4) and inter-subunit ratcheting (5), among other processes. The GTPase Elongation Factor G (EF-G) catalyzes this step via the consumption of GTP energy, which enhances the translocation kinetics by approximately 50,000-fold (6). Nonetheless, translocation can occur spontaneously without EF-G or in the presence of EF-G complexed with non-hydrolysable nucleotides (6–8). Therefore, it is not clear what is the essential role of energy in this process.

Structural studies have demonstrated that this 5-domain EF-G protein can undergo significant conformational changes, transitioning from highly compact to extended forms (9–13). After translocation, domain IV occupies the vacant A-site, which can function by either preventing the back-translocation of the A-site tRNA after its Brownian diffusion to the P-site or by actively promoting the translocation. The two proposed models, Brownian ratcheting, and power stroke, are not mutually exclusive, as they can proceed under different conditions, such as Mg^2+^ concentration and viscosity variations. Consequently, different experiments support either the combination of both models (12, 14, 15), the power stroke model (6, 16–18), or the Brownian ratchet model (8, 13, 19–22).

Regardless of the specific mechanism, the ribosome’s EF-G dependent movement on mRNA is essential for the proper functioning of the cell. For example, specific phosphorylation of human eukaryotic elongation factor 2 (eEF2) at residue Thr56 deactivates eEF2 and inhibits protein synthesis. Since the enzyme eEF2 kinase is highly specific in its role of mediating the phosphorylation on Thr56, it is a promising drug target for its ability to globally downregulate protein synthesis. Therefore, eEF2 kinase is a sought-after target in drug discovery research (23). On the other hand, the mechanism of deactivation on the Thr56 is less understood (24, 25).

In this report, we introduce T48E or T48V mutants on an engineered *E. coli* EF-G variant (named M5). The Thr48 residue, located in the switch I region of the GTP binding pocket, is highly conserved and sequence-aligned closely to Thr56 in human eEF2. Additionally, recent time-resolved cryo-EM studies show that switch I undergoes a large flipping motion (12, 13, 26). This motion creates a Pi release channel essential for EF-G’s function on the ribosome.

Consequently, several residues in this region could influence EF-G’s function, including Thr48. We monitor the interdomain changes in EF-G by measuring the distance between the residues 410 (located on the loop connecting domain II and III) and 533 (situated in domain IV). The compact and extended conformations are defined based on the relatively higher and lower FRET values generated by the dyes attached to these two residues (Figure 1). We show that these mutants trap the EF-G on the ribosome in a highly compact form prior to translocation, which is even more compact than free EF-G. In contrast, wild-type EF-G undergoes an extension on the ribosome upon GTP hydrolysis. This novel conformation may be due to T48 mutagenesis. We further discussed the potential biological significance of this observed conformation.

**Figure 1.**
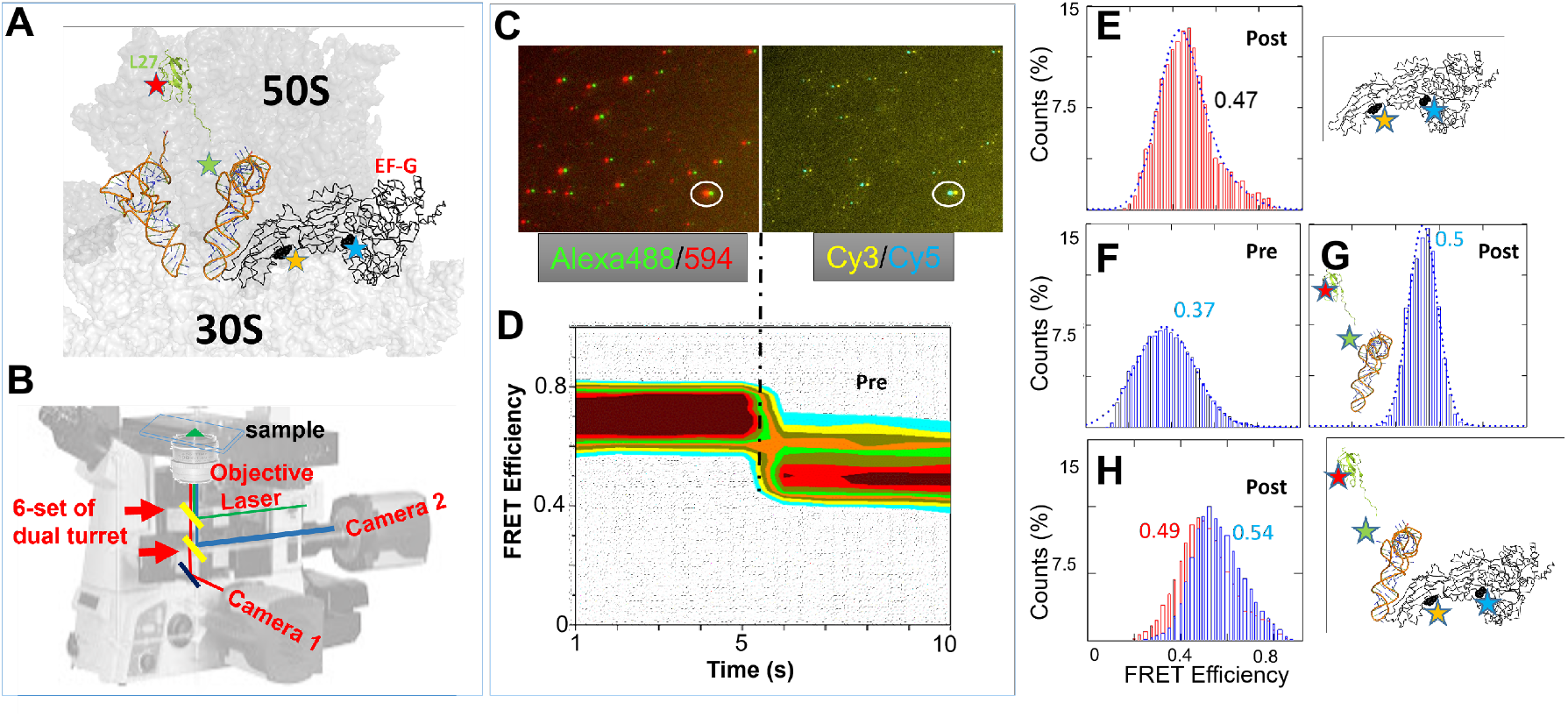
TIRF based smFRET instrument and representative data. Red histograms represent conformational changes within EF-G while blue histograms represent the changes between RbL27-tRNA. (A) Locations of the FRET pairs to simultaneously monitor ribosomal translocation and conformational changes in EF-G (positions of dyes are shown as stars in L27, tRNA^Phe^, and EF-G), and anticipated FRET diagrams for EF-G transition from pre- to post-translocation state. (B) Configuration of dual-turret/dual-camera TIRF microscope used in smFRET experiments. (C) An example of multi-channel dual camera imaging of Alexa488/Alexa594-labeled EF-G bound to the ribosome with Cy3-labeled Phe∼tRNA^Phe^ and Cy5-labeled L27 (D) 2D contour FRET efficiency diagram representing the imaged FRET state depicted in C.(E) FRET efficiency histogram of labeled EF-G in complex with unlabeled ribosome. (F-G) FRET efficiency histograms of Pre- and Post-translocation ribosomes depicted by L27-tRNA labeled ribosome complexes. (H) FRET efficiency histogram of both EF-G and Post-translocation ribosome in the same complex. The particle counts for the FRET efficiency histograms can be found in Table S1.

## MATERIALS AND METHODS

More details of the methods are in the supplementary file.

### Cys-EF-G mutagenesis and labeling

The Cysteineless 6xHis-tagged EF-G cloned in pET24b (27) was mutated with two Cysteine residues at the Phe410 and Tyr533 positions (*E. coli.* Sequence # without methionine (12)) via the “GeneArt Site-Directed Mutagenesis PLUS” kit, which is named M5 as previously reported (18). This mutated plasmid was transformed and expressed in BL21(DE3) pLysE cell, and purified with HisTrap^TM^ HP 5ml column on an Akta Purifier FPLC instrument with gradients (GE Healthcare Life Sciences). The clarified cell lysate was loaded on the column equilibrated with buffer A (50 mM Tris-HCl, pH 8, 50 mM NaCl), and eluted with a gradient from 5 to 500 mM imidazole. The His-tagged protein eluted around 200 mM imidazole. The protein was concentrated with Amicon Ultra-15 centrifugal filters (Millipore) and buffer exchanged to storage buffer (20 mM Tris-HCl, pH 7.5, 10 mM MgCl2, 0.5 mM EDTA, 4 mM BME, 400 mM KCl) using NAP-25 desalting column (Cytiva) (18). The protein was then labeled with Alexa 488- and 594-maleimides (ThermoFisher) following the manufacturer’s protocol. The final double labeling efficiency on M5 is approximately 60% (**Figure S1**).

### Ribosome labeling

The ribosome was isolated and labeled as described before (28). The *E. coli* strain IW312 lacking the gene *rpmA* encoding L27 was obtained from Dr. Robert Zimmermann of University of Massachusetts, Amherst, MA (29). Briefly, the IW312 ribosome was isolated, meanwhile, the L27 was expressed from the *rpmA* gene that was subcloned in the pET-20b(+) plasmid. The single natural Cys53 residue was labeled with Cy5-maleimide, after which it was incorporated into the ribosome. After this incorporation, the reconstituted ribosome was isolated from unbound L27 protein through the application of sucrose cushion centrifugation.

### Cy3-tRNA^Phe^ labeling

The yeast tRNA^Phe^ from Sigma was labeled on its dihydrouridine (D) residues at the 16/17 positions using Cy3-hydrazide from GE Healthcare. This process involved first reducing the D residues with sodium borohydride (NaBH4) at a neutral pH while maintaining the mixture on ice for 1 hour. Subsequently, the labeling with the dye was performed at a pH of 3.7 over a period of 2 hours at 37 °C, following a published procedure (30).

### Ribosome complex formation

All the mixtures were in TAM10 buffer, which consisted of 20 mM Tris-HCl (pH 7.5), 10 mM MgCl2, 30 mM NH4Cl, 70 mM KCl, 0.5 mM EDTA, and 7 mM BME (2-mercaptoethanol). Four mixtures were prepared (28): the ribosome mix, TuWG mix, Tu0G mix, Phe mix. The ribosome mix contains 1 μM ribosome, 1.5 μM each of initiation factors IF1, IF2, IF3, 2 μM of mRNA, 4 μM of charged fMet-tRNA^fMet^, and 4 mM of GTP. The TuWG mix contained 4 μM EF-Tu, 2 μM EF-G, 4 mM GTP, 4 mM 2-phosphoenolpyruvate (PEP), and 0.02 mg/mL Pyruvate Kinase. The Tu0G mix contained all the components found in the TuWG mix, with the singular exception of EF-G. The Phe mix contained 100 mM Tris (pH 7.8), 20 mM MgCl2, 1 mM EDTA, 4 mM ATP, 7 mM BME, 2 μM tRNA^Phe^ synthetase (PheRS), 2 A260 units/ml Cy3-labeled tRNA^Phe^ (30), and 0.25 mM phenylalanine. The ribosome mix, Tu0G mix, and Phe mix were combined in a volume ratio of 1:2:2, and then incubated at 37 °C for 2 minutes. The resulting MF-Pre ribosome complex was added on top of 1.1 M sucrose cushion and purified by centrifugation with a Hitachi CS150FNX ultracentrifuge (S140AT rotor, 200,000-400,000xg, 4℃, 3 hours). The post-translocation complex was prepared in a similar manner, except that the TuWG solution was used in place of the Tu0G solution. Additionally, the incubation time was extended to 10 minutes.

### SRL/ribosome and EF-G complexes for smFRET experiments

The 5’-biotinylated SRL (1 microM) is incubated with 1 microM EF-G•GTP in TAM10 buffer for 10 min at 37 °C. Then diluted to ∼ 10 nM and tethered to the surface according to **Figure S2**. In addition to double labeled EF-G experiments as showing in **Figure 5** of the main text, single Cy3-labeled EF-G is shown to FRET efficiently with Cy5-labeled SRL (Vector laboratories’ 3’-labeling Kit), indicating proper binding of these two molecules (FRET histogram in **Figure S2**).

The ribosome Cy3/Cy5 labeled MF-pre complex is assembled from three stocks, Initiation Complex (IC) Premix, Elongation Factor (EF) Premix and Aminoacylation (AM) Premix, prepared separately and combined at 1:2:2 volume ratio, respectively. All of the stocks were prepared using TAM10 buffer (1x TAM10: 20 mM Tris-HCl, pH 7.5, 10 mM MgAc2, 30 mM NH4Cl, 70 mM KCl, 0.5 mM EDTA, 7 mM BME (β-mercaptoethanol)). IC Premix contained 1.5x TAM10, 20 mM Tris-HCl, pH 7.5, 1 microM Biotin-RNA (coding sequence MFEKK…), 0.2 microM Cy5-L27-labeled ribosomes, 1.6 microM fMet-tRNA^fMet^, 1/2/1 microM of IF1/2/3, respectively, 1 mM GTP, and was preincubated at 37°C for 30 min. EF Premix contained 1x TAM10, 32 mM Tris-HCl, pH 7.5, 4 mM GTP, 4 mM PEP, 100 microg/ml pyruvate kinase, 12 microM EF-Tu, and was preincubated at 37°C for 30 min. AM Premix contained 0.8x TAM10, 33 mM Tris-HCl, pH 7.5, 4 mM ATP, 50 microM L-Phe, 2 A260 units/ml E.coli Cy3-labled tRNAPhe (charging capacity 800 pmol Phe/A260 unit), 10 ng/ml PheRS, and was preincubated at 37°C for 20 min. The three solutions were mixed and incubated at room temperature for 1 minute. The final MF-pre ribosome complex was layered on 1.1 M sucrose cushion and pelleted at 750,000xg and 4°C for 3 hours (Hitachi CS150FNX ultracentrifuge with S140AT rotor). Supernatant was carefully removed and pellet was resuspended in TAM10 buffer. 1 microM of this MF-pre complex was incubated at 37°C for 10 min with 1 micro EF-G, 1 mM of varied guanidine nucleotides, 1 mM fusidic acid. The complex was then diluted to 10-100 nM for smFRET imaging. The surface tethering strategy is shown in Figure S4.

### Single-molecule FRET imaging

The ribosome complexes were tethered to the biotinylated glass microscope coverslip at an appropriate concentration, ranging from 10 to 100 nM (**Figure S2**). This attachment was facilitated by the biotin-streptavidin-biotin interaction with biotinylated mRNA. An oxygen scavenger cocktail (0.8% glucose, 1 mg/mL glucose oxidase, 0.02 mg/mL catalase, and 2 mM Trolox) was added to the channel before imaging to prevent photobleaching (28). Single-molecule Förster Resonance Energy Transfer FRET movies were recorded using a dual-turret/dual camera Total Internal Reflection Fluorescence (TIRF) microscope, which is based on a Nikon Eclipse Ti2-E inverted microscope. This set-up utilizes a Coherent® OBIS™ 488 nm LS Laser and a LaserQuantum® solid state 532 nm Laser. The microscope is equipped with a top turret that reflects a laser for TIRF illumination, and a bottom turret that divides the FRET signal based on wavelength, directing it to two separate Complementary Metal Oxide Semiconductor (CMOS) cameras. Typically, 40 fields of view are gathered on a pre-programmed stage. In each field of view, a movie of 100 frames is recorded, with 100 ms exposure time. The fluorescence intensities of FRET pairs are fitted with ImageJ software and then analyzed with Mathcad program. The data analysis is detailed in supplementary file (**Figure S5**). The particle counts for all the FRET efficiency histograms can be found in Table S1.

## RESULTS

### Multi-Channel Single-Molecule FRET (smFRET) instrument

**Figure 1A** depicts the topology of EF-G, 30S/50S and tRNAs. The EF-G domain IV interacts with the A-site tRNA to accelerate the translocation, which is approximately 90 Å away from the GTP binding pocket. An allosteric mechanism must therefore be in effect to link the GTP hydrolysis with the conformational change occurring in domain IV of EF-G. To monitor the tRNA translocation from the A-site to the P-site in the ribosome, a FRET pair consisting of Cy5-L27 and Phe-tRNA^Phe^-Cy3 is incorporated into the pre-translocation ribosome complex, as illustrated in **Figure 1A**. As the tRNA progresses from the A-site to the P-site, the distance between the tRNA and L27 (a protein component of the 50S) decreases (28). This change in proximity leads to an increase in the FRET efficiency from 0.37 to 0.5, representing successful translocation (**Figures 1F-G**). Meanwhile, M5, the F410C-Y533C variant of a cysteineless *E. coli* EF-G (18), is labeled with a separate FRET pair, Alexa488-594 at positions 410/533 (**Figure 1A and E**, and in **Figure S4**). This FRET pair allows us to monitor conformational changes of EF-G, which are expected to occur concurrently with tRNA movement. This experimental setup thus presents a multi-channel, dual-FRET-pair-monitoring system for observing both tRNA movement and EF-G conformational changes on the same biomolecule complexes (**Figure 1H**). The single and dual FRET pair complexes exhibited consistent results, indicating fluorophore crosstalk is not a serious problem. The correction for the spectrum is detailed in **Figure S3**, along with its accompanying description.

The objective-based total internal reflection fluorescence (TIRF) microscope was constructed using a Nikon Eclipse Ti2-E inverted microscope equipped with two auto-turrets and two CMOS cameras (**Figure 1B**). The top turret houses filters that direct laser light towards the objective for Total-Internal-Reflection illumination, while the bottom turret contains filters that split the FRET fluorescence, directing the donor and acceptor emissions to the back and right cameras, respectively. By rotating the turret, up to six FRET pairs can be monitored simultaneously in a single sample, although two FRET pairs are demonstrated in this report. In one field of view, the microscope captures 50 images (100 ms/frame) with the 488-laser illumination and Alexa 488/495 filter cube, followed by 50 images with the 532-laser illumination and Cy3/Cy5 filter cube. **Figures 1C** and **S5** show one example of dual-FRET-pair-monitoring of the ribosome complex illustrated in **Figure 1A**. Fluorescent signals from the same physical location were superimposed in a combined movie of 10 seconds, even though the FRET pairs were on different parts of the complexes (EF-G, tRNA and ribosome). Consequently, the conformational changes of the EF-G and information about the tRNA translocation can be gathered from the same time-lapse trace, from the first and second half of the trace respectively. The FRET efficiency is calculated by the equation “FRET = (*I*_acceptor_)/(*I*_donor_+ *I*_acceptor_)” with spectrum corrections of crosstalk (**Figure S3**). The corresponding FRET efficiency histogram can then be rendered into a 2D contour format, effectively mapping the status of both the tRNA and the EF-G. For instance, **Figure 1D** presented such a contour representation of the FRET efficiency histogram. In this graph, the first 5-s represents FRET efficiency histogram of Alexa 488/594 pair. In addition, the FRET analysis is grouped by time points, to show its time-resolved potential. The second 5-s segment represents FRET efficiency histogram of Cy3/Cy5 FRET pair, also expanded with time-lapse. The X- and Y-axis depict the time and FRET efficiency bins, respectively, and the color depict the counts in histogram bins, similar to other contour plots (31).

This diagram indicated that in this specific ribosome complex, the EF-G was in a conformation characterized by a high FRET efficiency (approximately 0.7). Simultaneously, the tRNA has relocated to the P-site, as indicated by a FRET efficiency of approximately 0.5. The estimated distances between the labeled residues may not perfectly match the calculated distances from the FRET values because microenvironment, the orientation and fluctuations of linkers often result in considerably lower FRET efficiencies. Thus, in this report, we employ relative rather than absolute FRET values to distinguish between the compact and extended conformations of EF-G (**Figure S4**). In conclusion, our multi-channel dual-FRET-pair-monitoring strategy has enabled us to concurrently observe and analyze two processes within a single complex. The detail of single molecule data analysis is available in **Figure S5**.

### Translocation catalyzed by M5 with different nucleotides

Utilizing this multi-channel smFRET set up, we observed correlations between the EF-G conformational change and ribosomal translocation. The pre-translocation complex of the ribosome, with labeled Cy5-L27, carried the fMet-Phe-tRNA^Phe^-Cy3 at the A-site and a vacant tRNA^fMet^ at the P-site. This complex was then incubated with Alexa488/594 labeled EF-G that was bound with either GTP (**Figures 2A, 4D**), GDP (**Figures 2B, 2E**), or GDPCP (**Figures 2C, 2F**), to initiate the process of translocation. The state of the ribosome was evaluated by the FRET efficiencies between the tRNA and the L27 protein. After the labeled fMet-Phe-tRNA^Phe^-Cy3 moved from the A-to the P-site, post-translocation ribosome exhibited FRET efficiencies around 0.5. Conversely, pre-translocation ribosome displayed FRET efficiencies around 0.4 because the peptidyl-tRNA did not translocate. These ribosome subpopulations were sorted into two categories: **Figures 2A-C**, and **2D-F** correspond to pre- and post-ribosomes, respectively. The small differences in Gaussian centers are insignificant due to experimental errors, considering the sigma of the Gaussian function (The fitted parameters are displayed in **Table S2**). The ribosome status was depicted in the second half of the time-lapse traces. Subsequently, the conformation of the ribosome-bound EF-G was obtained from the first half of the same trace. These unique settings allowed us to establish correlations between the movements of EF-G and tRNA.

**Figure 2.**
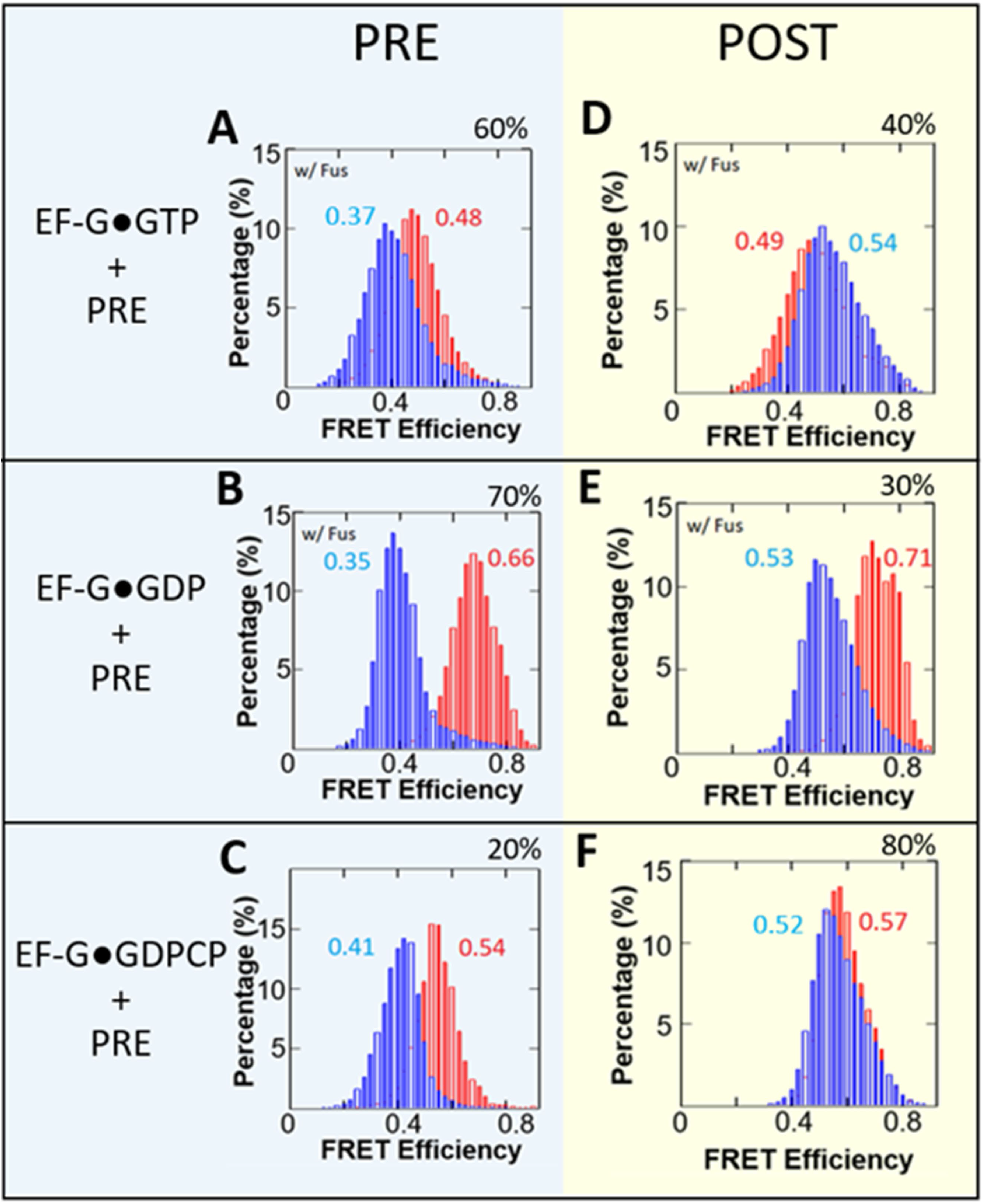
smFRET efficiency histograms for M5 EF-G catalyzed translocation. Red histograms represent conformational changes within EF-G while blue histograms represent the changes between RbL27-tRNA. Combined histograms are from a singular experiment. (A-C) FRET efficiency histograms for pre-translocation ribosome-bound with labeled EF-G. (D-F) FRET efficiency histograms for post-translocation ribosome-bound with labeled EF-G. (A,D) Experiments with EF-G•GTP; (B,E) Experiments with EF-G•GDP; (C,F) Experiments with EF-G•GDPCP. Experiments with EF-G•GTP and EF-G•GDP employed fusidic acid (Fus) to stabilize EF-G binding on the complexes. Ribosomal subpopulations of pre- or post-translocated ribosomes, expressed as a percentage, were obtained via sorting using predetermined FRET efficiencies for each state. The percentages are the ribosome translocation yields under EF-G in complex with the different nucleotides. The particle counts for the FRET efficiency histograms can be found in Table S1.

Under all nucleotide conditions, partial translocation was observed, with 40%, 30% and 80% completion for EF-G•GTP, EF-G•GDP, EF-G•GDPCP, respectively (**Figures 2D-F**). Figure 2 demonstrates that the double-labeled M5 can successfully translocate labeled tRNA from the A-site to the P-site in L27-labeld ribosome, which indicates its reasonable activity. In both the GTP and GDP experiments, fusidic acid (fus) was introduced to stabilize EF-G on the ribosome. As a result, both experiments showed reduced translocation efficiencies (under multiple turnover conditions) compared to the GDPCP condition, where no fusidic acid was used. While it has been reported that EF-G bound to GDP (EF-G•GDP) can promote translocation at a rate similar to that of a non-hydrolysable GTP analog, this is not the case when fusidic acid was present (31). Under these conditions, the efficiency of translocation promotion by EF-G•GDP was significantly reduced to 30% (**Figure 2E**).

After subpopulation sorting, the FRET efficiency histograms for EF-G and for tRNA-L27 pairs were extracted as illustrated in **Figure 1**. The EF-G conformations appeared to be largely similar regardless of whether the ribosome was in the pre- or post-state, and this held true for all three nucleotide forms (GTP, GDP, and GDPCP). This observation suggests that the conformation of EF-G may not have a direct correlation with the ribosomal state with or without fusidic acid, and agrees with a recent cryo-EM studies showing that EF-G acting more like a rigid body during translocation (12, 13). Fusidic acid (Fus) is known to prevent the release of EF-G from the ribosome after one round of GTP hydrolysis. Consequently, in **Figures 2A** and **D**, EF-G should have been trapped in the form of EF-G•GDP•Fus (32). On the other hand, **Figures 2B** and **E** showed outcome of EF-G•GDP directly interacting with the ribosome, in the presence of Fus. While the final chemical states were expected to be very similar, the conformation for the EF-G bound with GTP was much more extended than that of EF-G•GDP, which may suggest that the EF-G•GTP interacts with the ribosome with more flexibility, while EF-G•GDP is known to interact with ribosome poorly but nevertheless trapped on ribosome by Fusidic acid (16).

### Mutagenesis on EF-G switch I

After establishing the multi-channel FRET methodology, we studied the role of switch I. In the EF-G/eEF2 protein family, switch I is an important component of the GTP-binding domain. It facilitates a complex network of both inter- and intramolecular interactions, including closing the GTP binding pocket and contacts with both the large and small ribosomal subunits. In this study, M5 serves as the template for introducing the ’switch I’ mutants (18). These cysteines facilitate subsequent smFRET measurements. Specifically, the distance variations and corresponding changes in FRET efficiency values between these two residues serve as a measure of the extension of domain IV relative to other domains (**Figure S4**).

**Figures 3A-B** depicts a detailed view surrounding the GTP binding pocket of EF-G (For convenience, residues are numbered in the same manner as in the 7pjv/y structures) (12). Three loops: the phosphate-binding loop (P-loop, blue in **Figure 3B**), and switch I (red in **Figure 3B**) and II (magenta in **Figure 3B**), are located near the inorganic Pi molecule. His91 on switch II alters its position to trigger GTP hydrolysis, and the subsequent release of Pi initiates a significant conformational change in EF-G (11). A recent cryo-EM study showed that a refolding of this region was accompanied by the release of Pi (12). The amino acid residue Thr48 is located within this effector loop of switch I, which forms an H-bond with E27 near the P-loop. In addition, Thr48 is approximately 4.7 Å away from H366 in domain II. Replacing threonine with glutamic acid or valine will enable H-bonding between T48E and His366 or interrupt T48V and Glu27, respectively, which will potentially affect switch I’s refolding upon Pi release.

**Figure 3.**
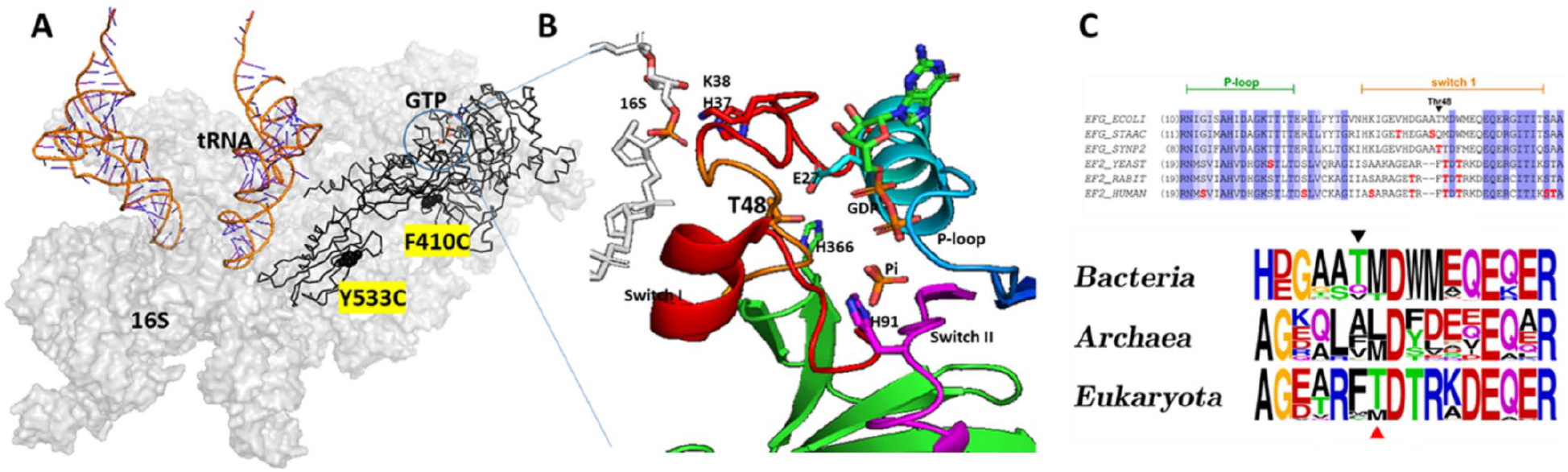
EF-G alignment, mutations, and smFRET labeling positions. (A) Topology of EF-G, tRNA and 16 S rRNA. The GTP binding site, and the two cysteines for smFRET experiments are highlighted. (B) Closeup view at the GTP binding pockets with some key elements and residues manifested. (C) Multiple sequence alignment of P-loop and switch I region and frequency sequence logo plots for a part of switch 1 region adjacent to the mutagenized threonine residue. Amino acid residues are shaded according to BLOSUM62 conservation score, darker shading or bigger font corresponds to higher conservation.

Additionally, T48 is close to residues H37/K38, which chelate with 16S rRNA. Therefore, T48 mutations could disrupt the sequential interaction between switch I and 16S rRNA during Pi release.

The alignment of sequences from *E. coli* EF-G and its analogous proteins in *S. cerevisiae* and *H. sapiens* (detail in supplemental material) reveals the conservation patterns in switch I and at the effector loop region (**Figure 3C**). As indicated in **Figure 3C**, at the position corresponding to Thr56 in human eEF2 (shown with a red arrow), threonine is predominantly found in eukaryotes, while methionine is more common in bacteria. However, a threonine residue, Thr48, is located nearby (indicated by a black arrow in **Figure 3C**). Although not reported for Thr 48 in *E. coli* EF-G, phosphorylation of threonine in the equivalent position was detected in EF-G from model cyanobacterium Synechococcus sp. PCC 7002 during global phosphoproteome analysis (32).

This observation suggests that bacterial EF-G might be subject to activity regulation through phosphorylation of amino acid residues in the switch I region, in a manner similar to their eukaryotic counterparts. Consequently, *E. coli* EF-G can serve as a model system to uncover the mechanism of Thr56-mediated inactivation in human eEF2. In our study, we introduced a phosphomimetic substitution at Thr48 with Glu (T48E). In addition, we created an auxiliary EF-G mutant, T48V, to assess the potential consequences of losing interactions attributed to the hydroxyl group of the original T48 residue.

### Enzymatic functional assays on T48V and T48E

To characterize the effect of T48V and T48E on EF-G function, we performed a series of in vitro functional assays (**Figure 4**). M5 and the T48E, T48V mutants have shown similar affinity in GTP/GDP binding (**Figures 4A**). The micromolar range binding affinities are consistent with literature values (33). The GTPase assay is using Transcreener GDP FI Assay system (BellBrooks Labs) in the end-point assay format (detail in supplemental information).(34) As shown in **Figure 4B**, the end-products of GDP are comparable in all three EF-G species. However, in contrast to the M5, T48V demonstrated a substantially diminished capacity to support poly(phenylalanine) synthesis on poly(U)-programmed Poly(Phe) assay. Moreover, the T48E variant exhibited almost no activity (**Figure 4C**). Co-sedimentation of EF-G variants with pre-translocation ribosomal complex in the presence of GTP were conducted and assessed via SDS-PAGE (**Figure 4D**). The gel revealed that T48V co-sedimented with the ribosomal S1 protein at approximately a 1:1 ratio, while T48E and M5 co-sedimented at 0.22:1 and 0.15:1 ratio, respectively. These findings suggest that the observed Poly(Phe) inhibition by T48V is due to impaired release from ribosome, while the inhibition by T48E appears to bind to the ribosomal pre-translocation complex and hydrolyzes GTP but leaves the ribosome without inducing translocation, or impairing in multiple round of translocation. It was shown that phosphorylation of eEF2 was unable to promote translocation as puromycin reactivity was not observed (24). This evidence might suggest that T48E is unable to catalyze translocation, similar to the effects of phosphorylation. The low co-sedimentation for M5 can be attributed to normal factor release following translocation.

**Figure 4.**
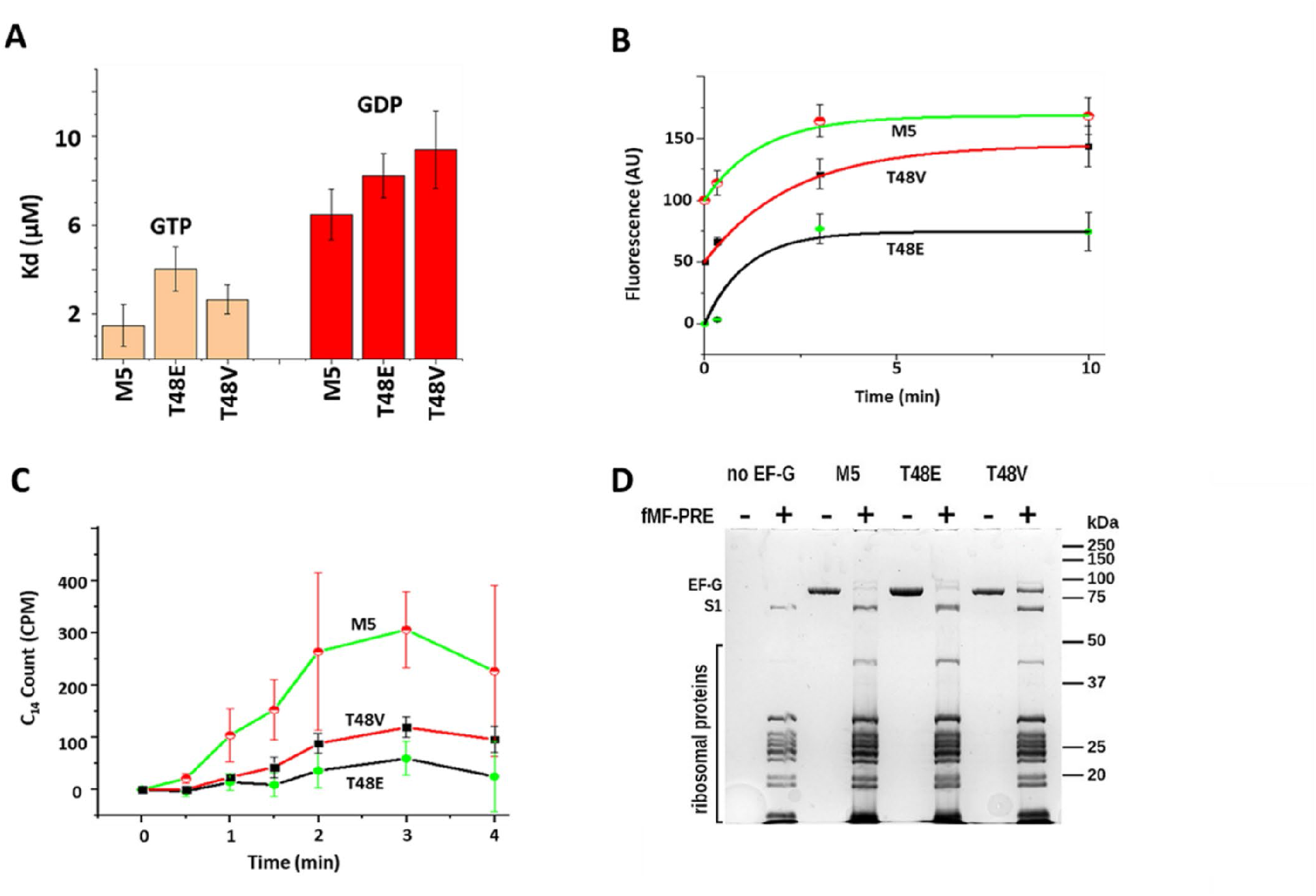
Functional assays on M5 and Thr48 mutated EF-G. (A) GTP/GDP dissociation constants of EF-G obtained by equilibrium fluorescent titration of EF-G variants with mant-GDP and mant-GTP at 20°C. (B) Ribosome-induced GTP hydrolysis monitored by a Transcreener GDP FI Assay system (BellBrooks Labs) at 20°C. Traces are shifted vertically by 50 units for clarity in display. (C) Kinetics of poly(U)-dependent poly([14C]Phe) synthesis at 37°C. The plots represent time-dependent synthesizing of [14C] labeled poly(Phe). (D) SDS-PAGE analysis of EF-G’s co-sedimentation with ribosomal pre-translocation complexes.

### Modulation of EF-G Conformation by the Ribosome

We further investigated the influence of the ribosome on the conformation of EF-G•GTP by comparing the conformation of free EF-G to EF-G when bound with the SR loop or the ribosome (Fus added with ribosome complexes). In their free form, both the M5 and T48E proteins adopted a similarly compact conformation (FRET value ∼ 0.7 in **Figures 5A** and **F**). On the other hand, the T48V mutant assumed a more extended conformation (FRET value ∼ 0.47 in **Figure 5K**). This suggests that the T48V mutant has caused a disturbance around the GTP binding pocket (**Figure 5B**). In the presence of sarcin-ricin loop (SRL), both M5 and T48E adopted more extended conformations (FRET value ∼ 0.47 in **Figures 5B** and **G**). However, the T48V mutant maintained its extended conformation, similar to its free form (**Figure 5L**) The change in FRET value from 0.7 to 0.45 suggests ∼ 10 Å extension of domain IV from domain III. The SRL is a highly conserved motif on the 23S rRNA and plays a crucial role in protein synthesis as it is the target of the highly toxic sarcin and ricin proteins. It is part of the ribosome GTPase activation center, promoting the GTPase activity in EF-G.(11–13, 35, 36) However, existing literature indicates that the SRL cannot independently trigger GTP hydrolysis without ribosome participation (37). Hence, **Figures 5B** and **G** suggest that the SRL likely prompted the extension of domain IV through its interaction with the interdomain region of I, III, and V in EF-G (**Figure S6**). Domains III and V serve as the two anchors maintaining the proximity of domain IV to the GTP binding domains. Therefore, the SRL might induce the extension of domain IV by loosening this interdomain region.

**Figure 5.**
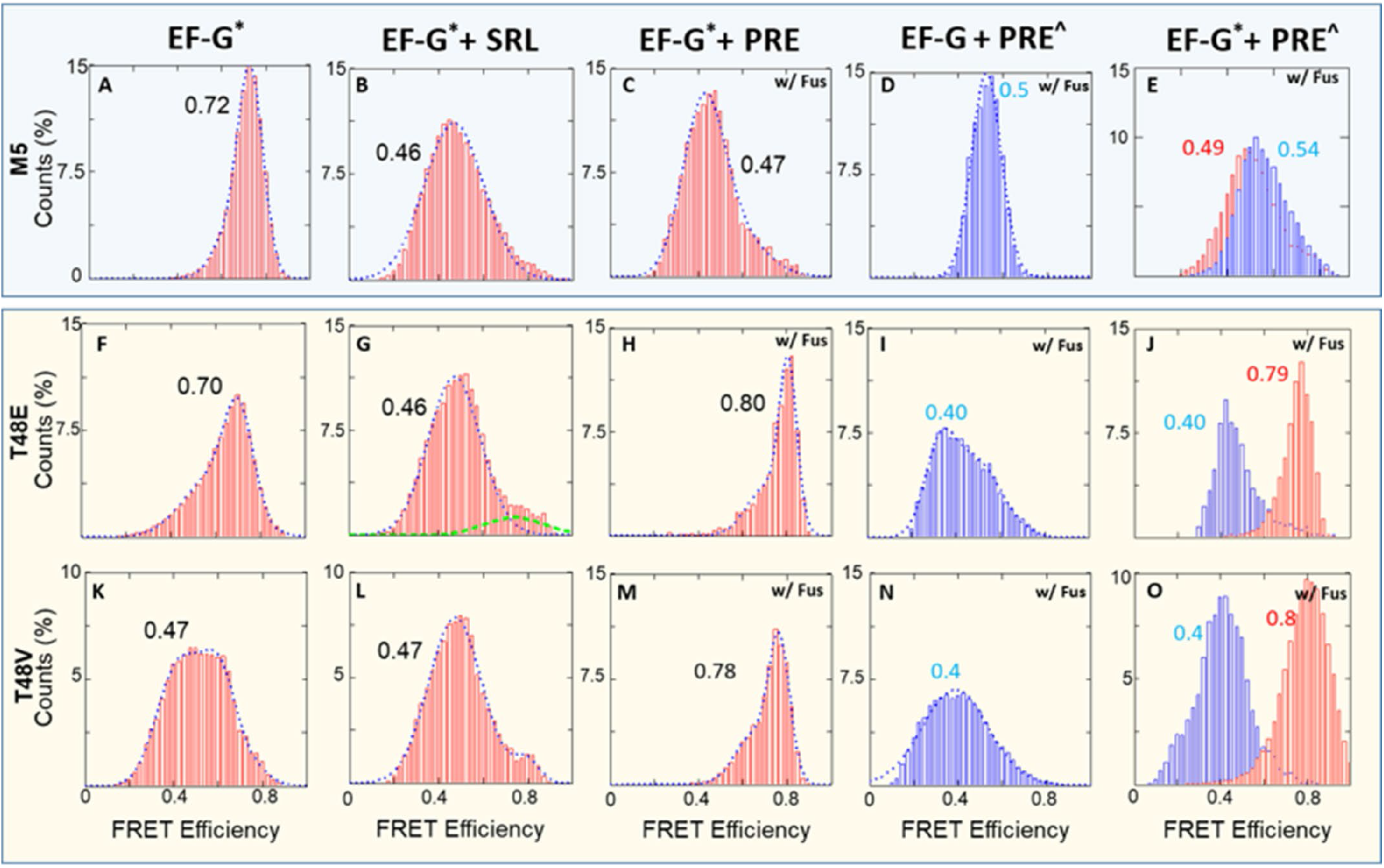
smFRET efficiency histograms for M5 and mutated EF-Gs. Red histograms represent conformational changes within EF-G while blue histograms represent the changes between RbL27-tRNA. (A-E) FRET histograms for EF-G variants M5•GTP in free form and bound with SRL, unlabeled pre-Rb, unlabeled EF-G/labeled pre-Rb, and labeled pre-Rb. (E-H) FRET histograms for EF-G variants T48E•GTP in free form and bound with SRL, unlabeled pre-Rb, unlabeled EF-G/labeled pre-Rb, and labeled pre-Rb. (I-L) FRET histograms for EF-G variants T48V•GTP in free form and bound with SRL, unlabeled pre-Rb, unlabeled EF-G/labeled pre-Rb, and labeled pre-Rb. The particle counts for the FRET efficiency histograms can be found in Table S1. * = labelling with Alexa 488/594; ^ = labelling with Cy3/Cy5.

When these EF-Gs interacted with pre-translocation complexes (without L27-tRNA labeling), **Figure 5C** demonstrated that M5 maintained a similar extended conformation as seen in **Figure 5B**. Conversely, the two mutant variants, T48E and T48V, transitioned into compact conformations, exhibiting a FRET efficiency between 0.7 and 0.8 (**Figures 5H** and **M**). This substantial difference in conformational adaptation was particularly remarkable when compared to the response observed in the case of SRL. These highly compacted EF-Gs were unable to promote translocation, as shown in **Figures 5I** and **N**, in which the EF-G were not labeled and only tRNA-L27 FRET pair was monitored. Finally, these single-FRET-pair observations were reproduced in the dual-FRET-pair results in **Figures 5E, J** and **O**. In these experiments, the pre- rb was labeled with L27-tRNA FRET pair to track the status of tRNA translocation, and EF-G was labeled with A488/A4594 pair. In contrast to the successful M5-assisted translocation depicted in **Figure 5E**, the two mutant variants failed to elicit the characteristic 0.5 FRET value observed in M5. Meanwhile, the L27-tRNA FRET pair demonstrated no translocation within these ribosome complexes (**Figures 5J** and **O**).

In summary, **Figure 5** suggests that the catalytic function of T48E and T48V EF-G was compromised. On the other hand, under M5 catalysis (Figure 5A-E), ribosomes bearing dual-FRET show the same FRET states as those with mono-FRET. This suggests minimal disturbance in activity due to labeling. An earlier study demonstrated that phosphorylated eEF2 cannot promote puromycin activity in a poly(U) assay, suggesting an impairment in the translocation step (24), which is consistent with our observation. In addition, ribosomal components beyond the SRL play a crucial role in orchestrating the allosteric transformation of EF-G, which occurs from its GTP binding site to the tip of domain IV 90Å away. In this study, we prepared the ribosome complexes in tubes prior to smFRET imaging. This approach was taken to enhance signal density and reduce negative surface effects on ribosomal activity. As a result, the single-molecule traces we obtained represent snapshots of ribosome complexes stabilized by fusidic acid. Kinetic information, derived from real-time delivery and signal acquisition, will be published in future studies.

## DISCUSSION AND CONCLUSIONS

We have developed a multi-channel dual-FRET-pair-monitoring experiment to detect the EF-G conformational change and the tRNA translocation in the same ribosome complexes. In addition to M5 EF-G, two mutations (T48E and T48V) are introduced, which may correspond to human eEF2’s Thr56 position. Phosphorylation at this position enables global downregulation of protein synthesis in human cells. The mechanism for this deactivation remains unclear, even though regulating of the eEF2 kinase, which phosphorylates Thr56, is of significant interest for the treatment of various diseases (38–40). We have found that both SRL and ribosome binding can stabilize EF-G•GTP at an extended form. However, the ribosome binding of these mutated EF-Gs results in a more compact, stable form that is inactive in translocation. This compact form is not observed in M5. Most structural studies revealed that EF-G, when bound to the ribosome, tends to adopt a more elongated conformation compared to its free form. **Figure S7** shows that when domain I is aligned, domain IV extends further from domain III, where the FRET-paired dyes are located. Notably, structure 3j9z adopts an intermediate state due to its H91A mutation, which impairs the GTP hydrolysis (11). On the other hand, the majority of observed structures exhibit a similar elongated conformation, regardless of the tRNA and nucleotide configurations, some of them also compared in **Figure S7**. Our results, while not providing exact measurements of distance changes, nonetheless reveal a FRET state with a higher FRET value compared to free EF-G (1fnm), indicating a more compact form than its free state. There is a reported example of EF-G bound to the ribosome with more compact from than free EF-G, likely due to artificial modification in the L9-fused EF-G preparation (9). However, the compact form we observed cannot be confidently assigned to this conformation. Despite this, it is still reasonable to assume the existence of other intermediate conformations between the free and ribosome-bound forms of EF-G. What we have observed can be novel conformation that is previously unobserved, potentially resulting from impairments in the GTP binding pocket.

In addition, an earlier FRET study have described similar compact conformations for the ribosome-bound, non-mutated *E. coli* EF-G. In these instances, these conformations were observed at low frequencies, contrasting with the predominantly extended conformation (41). Therefore, the results indicate that these compact conformations may occur naturally, but it is increased by the specific mutations.

The molecular mechanisms and interactions altered by the mutations can be better understood through the structural information provided in recent studies. **Figure S8** illustrates the differences in the GTP-binding pocket upon Pi release (12). Before Pi release (**Figure S8A**), the switch I encloses the GTP binding pocket. The significant downward swing of the effector loop, where the mutations are situated, opens the pocket (**Figure S8B**). This shift creates a visible corridor for the inorganic phosphate (Pi) to escape. Concurrently, several motifs from domain I (blue and light blue), II (green), III (yellow), V (salmon) move away from the GTP center (**Figures S8A** and **B**). **Figures S8C** and **D** illustrate the switch I region’s large swing displacement in ribbon models. In addition, interactions further away from the GTP also changed (**Figures S8E** and **F**). For instance, novel interactions can be observed between Thr325 and Glu440, Arp687/Glu688 and Arg494, as well as Met462 and Thr667. On the other hand, some interactions are weakened, such as those between Arg160 and Arg676, and between Met460 and Arg670. Overall, the interactions between domain III and V, as well as domain V to IV, are strengthened, while interactions between domain I and V are weakened. These changes enable domain IV to extend further away from the core structure of EF-G.

The mutation of threonine at position 48 to glutamic acid potentially creates a stronger interaction between the effector loop and residues His366/Lys369 in domain II. This can restrict the mobility of this element and inhibit the major swinging displacement that is associated with translocation. Conversely, the T48V mutant appears to have difficulty dissociating from the ribosome in the co-sedimentation results of **Figure 4D**, maybe due to new interactions via the hydrophobic valine residue. In both mutants, a more compact EF-G conformation is observed to be fixed on the ribosome in smFRET experiments.

Since GTP hydrolysis occurs normally and is responsive to ribosome enhancement (**Figure 4**), it is reasonable to assume that this highly compact conformation is trapped after GTP hydrolysis but is unable to induce translocation. Time-resoled cryo-EM visualization has revealed three intermediates of EF-G bound to ribosome, predominantly exhibiting extended form with small scale interdomain rearrangement. In these three states, the ribosome transitions from fully rotated state to a novel, previously unseen state with highly swivel head and further translocated tRNAs. Therefore, it is probable that the compact form we observed occurs before the states in cryo-EM. This is because it appears improbable that an additional compact state would occur amidst these states, which are shown to facilitate smooth ribosomal transitions (13). Thus, it is possible that this notably compact conformation represents a transient intermediate state resulting from the interactions mentioned earlier (**Figure S9)**. Nevertheless, it remains plausible that this condensed state is exclusively induced by specific mutations at position 48. Additional research is essential to uncover its biological significance.

## AUTHOR INFORMATION

### Corresponding Author

*

### Author Contributions

J.J., J.S., and R.L. have conducted the single molecule experiments; V.S. and M.G. conducted the mutagenesis and biochemistry assays. The manuscript was written through contributions of all authors. All authors have given approval to the final version of the manuscript.

## ACKNOWLEDGMENT

This research is supported by the US National Institutes of Health (R01GM111452), NSF (2130427) and the Welch Foundation (E-1721).

We thank the generous large equipment fund from the University of Houston. J.J. and M.G. are supported by the “HAMBP” (T32GM008280) predoctoral fellowship. J.S. is supported by NIH (3R01GM111452-07S1) predoctoral fellowship.

## PROTEIN ACCESS ID

The uniprot ID for *E. coli* EF-G is P0A6M8.

## Supporting information for

This supplemental material contains materials and preparation details, 2 supplement table, and 15 supplemental figures.

## MATERIALS AND PREPARATIONS

### Protein mutagenesis and expression

BL21 Star (DE3)pLysS and BL21(DE3)pLysE competent cells, and GeneArt Site-directed Mutagenesis PLUS Kit were purchased from ThermoFisher Scientific. Q5 Site-directed Mutagenesis Kit was purchased from New England Biolabs. The HisTrap^TM^ HP 5 ml column and Nap^TM^ desalting columns were acquired from Cytiva. Amicon Ultra centrifugal filters are from Millipore.

### Sarcin-ricin loop

RNA oligo of 29 nt and biotinylated at the 5’-end (5’-biotin-TEG-CUGCUCCUAG UACGAGAGGA CCGGAGUGG) was purchased from Integrated DNA Technologies.

### Fluorescent Dyes and mant-nucleotide

Alexa Fluor^TM^ 488 C_5_ maleimide (Alexa 488) and Alexa Fluor^TM^ 594 C_5_ maleimde (Alexa 594) fluorescent dyes were purchased from ThermoFisher Scientific, Cyanine3 maleimide (Cy3) and Cyanine5 maleimide (Cy5) fluorescent dyes were purchased from Lumiprobe. 2’/3’-O-(N-Methyl-anthraniloyl)-guanosine-5’-triphosphate (mant-GTP) and 2’/3’-O-(N-Methyl-anthraniloyl)-guanosine-5’-diphosphate (mant-GDP) were from Sigma Aldrich.

All the other materials were from Millipore-Sigma.

### mRNA preparation

The DNA template oligos for the *in vitro* T7 transcription reaction were purchased from Integrated DNA Technologies. The HiScribe T7 High-Yield RNA Kit was from New England Biolabs. The 70-nt mRNA (5’-GGGCAACUGU UAAUUAAAUU AAAUUAAAAA GGAAAUAAAA AUGUUUAAAC GUAAAUCUAC UGCUGAACUC-3’) with a ribosome-binding site and a 10-codon-long non-stop coding sequence for Met-Phe-Lys-Arg-Lys-Ser-Thr-Ala-Glu-Leu peptide was synthesized and purified according to manufacturer’s instructions.

### Ribosome preparation

The 70S ribosomes were purified from *E. coli* MRE600 according as previously described.(1) A culture of *E. coli* MRE600 was propagated in LB medium in a shaker incubator at 37°C and 200 rpm until optical density at 600 nm reached value of 0.6. The cells were harvested by centrifugation at 3,000xg, 4°C, for 20 min. The cell pellet was washed with cold buffer I (50 mM Tris-HCl, pH 7.6, 10 mM MgCl_2_, 100 mM NH_4_Cl, 6 mM BME, 0.5 mM EDTA). Then the cells were resuspended in the same buffer with 0.2 mg/ml egg white lysozyme and 2 μg/ml DNase I and incubated for 30 min on ice. Afterward the cells were lysed by sonication for 5 min (10 sec pulse, 20 sec pause) on ice. Cell debris were removed by two consecutive centrifugations at 14,500xg, 4°C, for 1 hour each. Cleared supernatant was layered on top of pre-cooled 1.1 M sucrose cushion in buffer II (20 mM Tris-HCl, pH 7.6, 10 mM MgCl_2_, 500 mM NH_4_Cl, 6 mM BME, 0.5 mM EDTA) with volume ratio 1:1, respectively. The ribosomes were pelleted by centrifugation on Beckman XL-80 ultracentrifuge with Type 45 Ti Fixed-Angle rotor at 120,000xg, 4°C, for 20 hours. The collected pellet was carefully rinsed with buffer I, then the ribosomes were resuspended in the same buffer, and the concentration of NH_4_Cl was adjusted to 400 mM. The ribosomes were pelleted again by ultracentrifugation at 120,000xg, 4°C, for 20 hours. The pellet was rinsed with buffer I, and finally resuspended in minimum volume of the same buffer. The ribosome concentration was determined by UV absorbance measurement at 260 nm using conversion coefficient 1 A_260_ unit = 23 pmoles of 70S ribosomes. The prepared ribosome solution was aliquoted, flash-frozen in liquid nitrogen, and stored at -80°C.

### Ribosome factors and aminoacyl-tRNA synthetases preparation

*E. coli* His-tagged IF1, IF2, IF3, EF-Tu, EF-G, methionyl- and phenylalanyl-tRNA synthetases were expressed and purified as previously described.(1, 2)

### fMet-tRNA^fMet^ preparation

*E. coli* tRNA^fMet^ was overexpressed in BL21 Star (DE3)pLysS cells from a recombinant gene cassette controlled by lpp promoter on pBluescript II SK (+) plasmid (Genscript). After cell lysis by sonication, tRNA^fMet^ was purified on Sepharose 4B column eluted with the reverse gradient of Ammonium Sulfate, and had methionine-accepting activity of 500 pmol/A_260_ unit. The recombinant 6xHis-tagged *E. coli* methionyl-tRNA synthetase and *E. coli* methionyl-tRNA^fMet^ formyltransferase were expressed in BL21 Star (DE3)pLysS cells and purified on HisTrap^TM^ HP 5 ml column according to manufacturer’s protocol. The formyl donor, 10-formyltetrahydrofolate, was prepared according to literature.(3) Aminoacylation of tRNA^fMet^ with subsequent formylation of Met-tRNA^fMet^ was performed as a one-pot reaction in a mixture containing 100 mM Tris-HCl (pH 7.6), 4 mM ATP, 20 mM MgCl_2_, 10 mM KCl, 150 μM L-Methionine, 750 μM neutralized formyl donor, 7 mM BME, 20 μM tRNA^fMet^, 12 μM methionyl-tRNA synthetase, and 16 μM methionyl-tRNA^fMet^ formyltransferase. The reaction mixture was incubated at 37°C for 1 hour, then acidified with sodium acetate (pH 5.0) added to the final concentration of 0.5 M. fMet-tRNA^fMet^ was purified by successive phenol and chloroform extractions followed by gel-filtration on NAP-10 column (Cytiva) equilibrated with 0.2 M sodium acetate (pH 5.0). Finally, fMet-tRNA^fMet^ was precipitated with ethanol, the precipitate was washed with 70% (v/v) ethanol, air-dried, and dissolved in 2 mM sodium acetate (pH 5.0). The solution was stored at -80°C.

### N-Ac-Phe-tRNA^Phe^_yeast_ preparation

N-Acetyl-Phe-tRNA^Phe^ was prepared from yeast tRNA^Phe^ (Millipore-Sigma, 1000 pmol/A_260_ unit) according to described procedure.(4)

### poly(Phe) synthesis assay

poly(U)-dependent synthesis of poly(L-[^14^C]Phe) was performed in a reaction mixture assembled from three stocks, Initiation Complex (IC) Premix, Elongation Factor (EF) Premix and Aminoacylation (AM) Premix, prepared separately and combined at 1:2:2 volume ratio, respectively. All of the stocks were prepared using TAM_10_ buffer (1x TAM_10_: 20 mM Tris-HCl, pH 7.5, 10 mM MgAc_2_, 30 mM NH_4_Cl, 70 mM KCl, 0.5 mM EDTA, 7 mM BME (β-mercaptoethanol)). IC Premix contained 1.5x TAM_10_, 20 mM Tris-HCl, pH 7.5, 2 mg/ml poly-U RNA, 0.4 μM ribosomes, 1.2 μM N-Ac-Phe-tRNA^Phe^_yeast_, and was preincubated at 37°C for 30 min. EF Premix contained 1x TAM_10_, 32 mM Tris-HCl, pH 7.5, 4 mM GTP, 4 mM PEP, 100 μg/ml pyruvate kinase, 12 μM EF-Tu, 4 μM EF-G (or no EF-G in a control), and was preincubated at 37°C for 30 min. AM Premix contained 0.8x TAM_10_, 33 mM Tris-HCl, pH 7.5, 4 mM ATP, 50 μM L-[^14^C]Phe, 120 A_260_ units/ml *E.coli* total tRNA (charging capacity 180 pmol Phe/A_260_ unit), 10 ng/ml PheRS, and was preincubated at 37°C for 20 min. AM Premix was combined with EF Premix and incubated at 37°C for 5 min, then IC Premix was added to initiate poly(L-[^14^C]Phe) synthesis. At appropriate time intervals, 12 μl aliquots were withdrawn from the reaction mixture, quenched with 6 μl of stop solution (1 M KOH, 25 mM EDTA), incubated at 37°C for 1 hour to destroy unused [^14^C]Phe-tRNA, and acidified with 6 μl 1.5 M sodium acetate (pH 4.5). Finally, 20 μl from each processed aliquot was applied onto a Whatman paper filter pre-impregnated with trichloroacetic acid (TCA). The filters were batch-washed with one change of 10% TCA, five changes of ice-cold 5% TCA, one change of ice-cold ethanol, and air-dried at room temperature for 30 min. Radioactivity of the filters was determined in ScintLogic U scintillation liquid (LabLogic) using Hidex 300 SL liquid scintillation counter.

### Equilibrium titrations of EF-G with mant-GTP and mant-GDP

Experiments were performed at room temperature (20°C) with samples prepared in Dilution buffer (50 mM Tris-HCl, pH 7.5, 70 mM NH_4_Cl, 30 mM KCl, 10 mM MgCl_2_, 0.5 mM EDTA, 6 mM BME). A titration grid of EF-G (M5, T48E or T48V) and mant nucleotide (mant-GTP or mant-GDP) was created with concentrations of 0, 2, 4, 8, 16, 32 μM EF-G and 0, 2, 4, 8, 16 μM mant nucleotide, respectively. Fluorescence was measured using a DeNovix QFX fluorometer set with excitation wavelengths in 361-389 nm range and fluorescence detection in 435-485 nm range. For each combination of an EF-G variant and mant nucleotide the titration grid was triplicated, and the fluorescence intensities from the three replicates were averaged. The obtained values were fitted to the following two-variable equation:

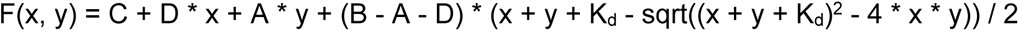

where F(x, y) is the fluorescence intensity in relative fluorescence units (RFU), x and y are the concentrations of EF-G and mant nucleotide, respectively, in μM, A is the specific molar fluorescence of free mant nucleotide in RFU/μM, B is the specific molar fluorescence of EF-G*mant nucleotide complex in RFU/μM, C is the background fluorescence in the absence of EF-G and mant nucleotide in RFU, D is the specific molar fluorescence of free EF-G in RFU/μM, and K_d_ is the dissociation constant for EF-G*mant nucleotide complex in μM.(5) The titration curves are displayed in Figures S10-S15.

### Ribosome-induced GTPase activity of EF-G variants

GTPase activity of EF-G variants was measured using Transcreener GDP FI Assay system (BellBrooks Labs) in the end-point assay format.(6) The assay is based on quantification of fluorescent GDP-derived tracer released from its complex with GDP-specific antibody conjugated with a quencher. Upon GTP hydrolysis by EF-G, the produced GDP displaces the tracer from the complex, which causes the increase of fluorescence emission intensity due to de-quenching. GTPase reaction mixture contained 0.75 μM EF-G, 0.125 μM *E. coli* ribosomes, 100 μM GTP, 20 mM Tris-HCl, pH 7.5, 10 mM MgCl_2_, 40 mM KCl, 0.5 mM EDTA, 4 mM BME. The reaction was initiated by addition of GTP, carried out at 37°C for several time intervals, and stopped by mixing with equal volume of the Stop & Detect solution containing 90 μg/mL GDP-specific antibody and 4 nM Alexa Fluor^TM^ 594-derivatized GDP tracer in 10 mM HEPES-NaOH, 20 mM EDTA, 0.01% Brij-35, pH 7.5. Afterwards, the mixture was incubated at room temperature (20°C) for 1 hour in the dark to allow equilibration between the antibody, tracer, and GDP. The fluorescence intensity was measured using DeNovix QFX Fluorometer with excitation wavelength range between 490-558 nm and emission detection in 560-650 nm range. The background GDP concentration was accounted for by running GTPase reaction in the absence of ribosomes while keeping all other conditions same. Each measurement conditions was repeated at least three times.

### EF-G/ribosome co-sedimentation assay

The pre-translocation ribosome complex carrying fMet-Phenyl-tRNA^Phe^ at the A-site and vacant tRNA^fmet^ at the P-site (fMF-pre, 0.4 μM) was incubated at 37°C for 5 min with 4 mM GTP and 2 μM of either M5, or T48E, or T48V variant of EF-G in TAM_10_ buffer. Afterwards, the concentration of Mg^2+^ in the binding mixtures was adjusted from 10 to 30 mM. The samples were placed on ice, then layered on top of pre-cooled 1.1 M sucrose cushions in TAM_30_. The ribosomal complexes were pelleted through the sucrose cushion at 750,000xg and 4°C for 3 hours (Hitachi CS150FNX ultracentrifuge with S140AT rotor). Supernatants were carefully removed; each pellet was resuspended in 10 μl of TAM_10_ buffer and resolved in a 10% SDS PAGE gel (200V/1hr). The gel was stained with Coomassie Brilliant Blue R-250, destained with 20% (v/v) ethanol, 5% (v/v) acetic acid, and photographed using ChemiDoc Touch Imaging System (Bio-Rad).

### Multiple sequence alignment of P-loop/switch 1 regions

Several representative EF-G/EF2 proteins are aligned: EFG_ECOLI, EF-G from E. coli strain K-12; EFG_STAAC, EF-G from Staphylococcus aureus strain COL; EFG_SYNP2, EF-G from Synechococcus sp. strain ATCC 27264 / PCC 7002 / PR-6 (Agmenellum quadruplicatum); EF2_YEAST, eEF2 from Saccharomyces cerevisiae strain ATCC 204508 / S288c; EF2_RABIT, eEF2 from Oryctolagus cuniculus; EF2_HUMAN, eEF2 from Homo sapiens. Position numbering of EFG_ECOLI, EF2_YEAST, EF2_RABIT, and EF2_HUMAN corresponds to the mature protein sequences with removed N-terminal methionine residue. Position numbering of EFG_STAAC and EFG_SYNP2 is derived from the corresponding gene sequences with retention of the N-terminal methionine. The sequences were aligned using SeaView software and further refined with structurally validated corrections. Boundaries of the P-loop and switch 1 are given according to Pulk A et al.(7)

**Table S1.** Particle counts in FRET efficiency histograms.

|  |  |  |  |  |  |  |
| --- | --- | --- | --- | --- | --- | --- |
| Figure ID | 1E | 1F | 1G | 1H |  |  |
| Particle counts | 1015 | 4779 | 3161 | 1827 |  |  |
| Figure ID | 2A | 2D | 2B | 2E | 2C | 2F |
| Particle counts | 731 | 1827 | 2779 | 1191 | 461 | 1843 |
| Figure ID | 5A | 5B | 5C | 5D | 5E |  |
| Particle counts | 1930 | 1201 | 1015 | 3161 | 1827 |  |
| Figure ID | 5F | 5G | 5H | 5I | 5J |  |
| Particle counts | 8913 | 1491 | 3078 | 3176 | 2328 |  |
| Figure ID | 5K | 5L | 5M | 5N | 5O |  |
| Particle counts | 889 | 720 | 885 | 5929 | 2326 |  |
| Figure ID | S2 |  |  |  |  |  |
| Particle counts | 626 |  |  |  |  |  |

**Table S2.** Fitted Gaussian positions (xo) and standard deviations (σ) for Figure 2. Values were calculated using the equation f(x)=Ae^(-〖(x-x_o)〗 ^2/〖2σ〗 ^2) in MathCad. All values were fit using a singular fitting.

| | L27-tRNA $x_0$ | L27-tRNA $\sigma$ | EF-G $x_0$ | EF-G $\sigma$ |
| --- | --- | --- | --- | --- |
| GTP pre-complex | 0.366 | $\pm 0.093$ | 0.483 | $\pm 0.085$ |
| GDP pre-complex | 0.347 | $\pm 0.077$ | 0.657 | $\pm 0.080$ |
| GDPCP pre-complex | 0.408 | $\pm 0.069$ | 0.540 | $\pm 0.065$ |
| GTP post-complex | 0.544 | $\pm 0.105$ | 0.490 | $\pm 0.111$ |
| GDP post-complex | 0.533 | $\pm 0.086$ | 0.713 | $\pm 0.079$ |
| GDPCP post-complex | 0.520 | $\pm 0.086$ | 0.573 | $\pm 0.074$ |

**Figure S1.**
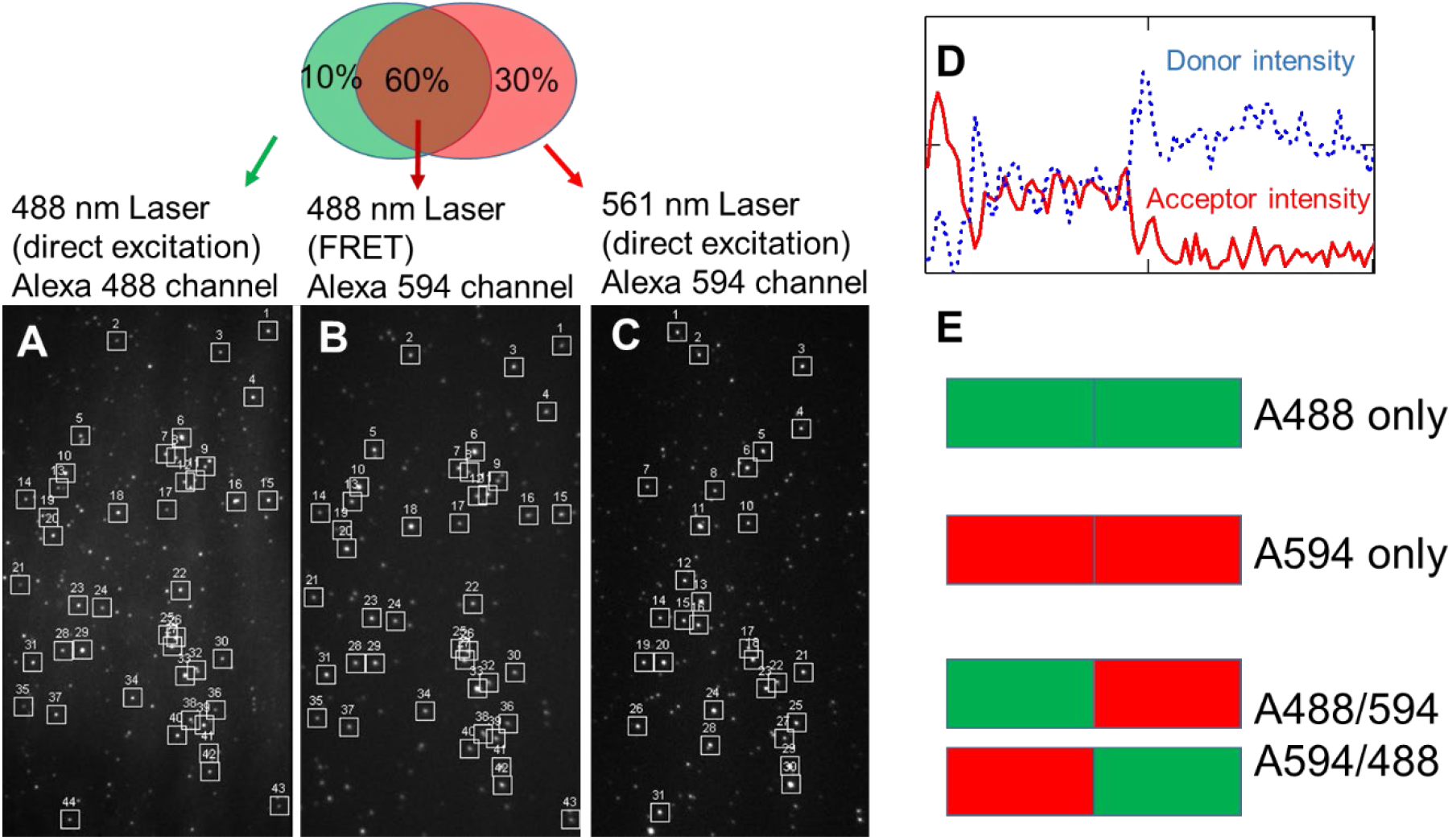
Analysis of Labeling composition of EF-G. **(A-B)** One image frame taken from a single movie showing directly excited Alexa 488 dye and FRET-excited Alexa 594 via a 488 nm laser, respectively. **(C)** One frame taken from a single movie showing directly excited Alexa 594 dye via a 561 nm laser. All images in A-C are from the same field of view. Squares show FRET pairs with ImageJ program. **(D)** Example of fluorescence emission intensities of the Alexa 488 (blue trace) and the Alexa 594 (red trace) over time. **(E)** Possibilities of the two cysteine labeling sites: both Alexa 488, both Alexa 594, or a combination of the two dyes. About 10% of labeling sites are Alexa 488 double labeled, 60% are a combination of Alexa 488/594, and 30% are Alexa 594 double labeled (detail of the analysis is given below).

The two unique cysteines at positions F410C and Y533C were simultaneously labeled with mono-maleimide Alexa Fluor^TM^ 488 (A488) and 594 (A594). Assuming equal labeling efficiency on both residues, three labeling compositions were possible: both A488, both A594, or (A488/594 or A594/488) (**Fig. S1E**). The compositions were measured via smFRET- and direct-excitations of the acceptor (A594). In the smFRET-excitation experiments, the 488 nm laser excited the A488 directly, which emitted in the A488 channel (**Fig. S1A**). Meanwhile, A594 emitted in the A594 channel via FRET (**Fig. S1B**) only if it was co-labeled on the same EF-G. The FRET pairs were fit with imageJ program (**Fig. S1C**). Therefore, the numbers of emitters in A488 and A594 channels corresponded to numbers of EF-G labeled with at least one A488 and double labeled with A488/594, respectively. Similarly, the number of EF-G labeled with at least one A594 was measured by emissions in the A594 channel under 560 nm laser (**Fig. S1C**), which only excited the A594 but not A488 dye. Consequently, the labeling composition can be represented as the overlapped ovals in the plot, in which the green and red ovals represented the EF-G being labeled with at least one Cy3 and Cy5, respectively; and the overlapped area represented the double labeled EF-G. The percentage of the EF-G labeled with A488/594 or A594/488 on C410/C534, respectively, was estimated to be approximately 60% (±7%, average of 300 field of views). The FRET efficiency histogram did not exhibit two peaks (**Fig. S3A**). Therefore, the mixture of double labeled EF-G (A488/A594 or A594/A488) was treated as one species. Other labeling components did not interfere with the FRET-data analysis because A594-only labeled EF-G did not generate any signal, and in the meantime A488-only labeled EF-G was not picked by the FRET-pair seeking program.

**Figure S2.**
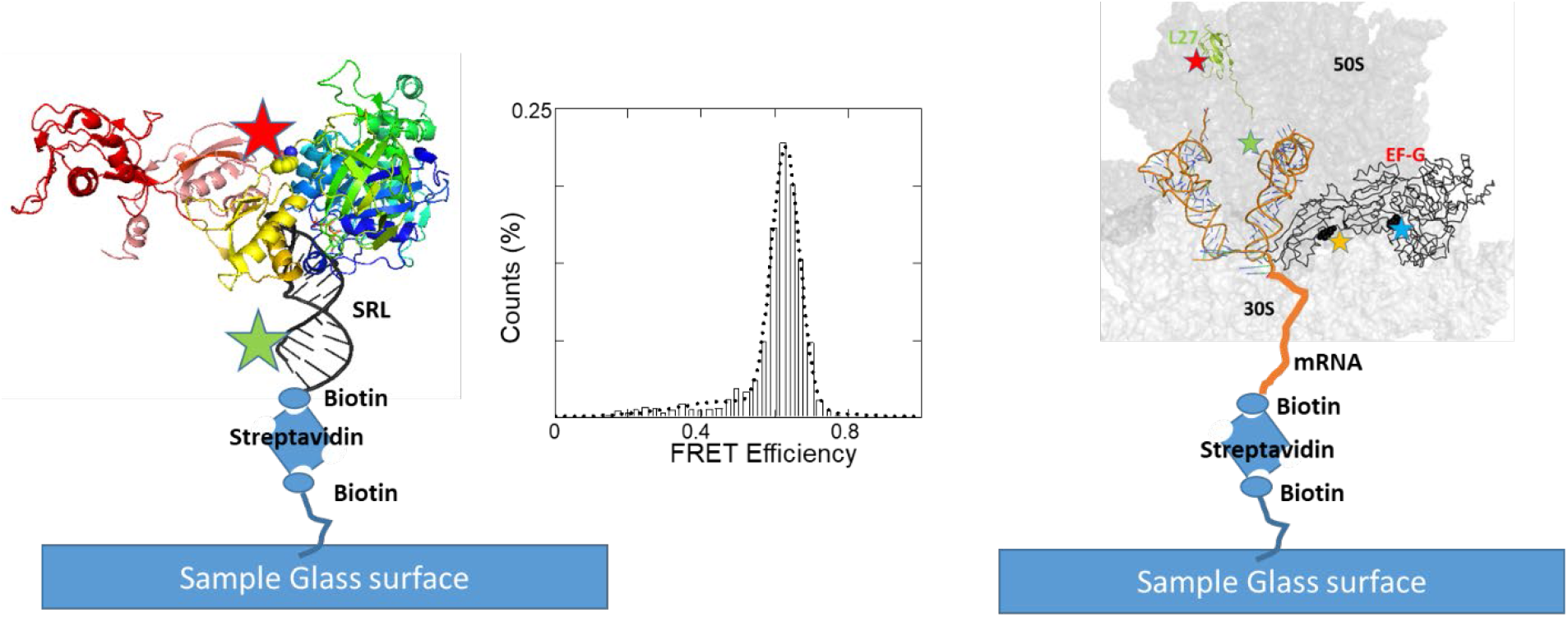
Surface tethering and FRET labeling strategies for SRL/Ribosome – EF-G complexes. The FRET efficiency histogram shows close proximity of 5’-biotin-SRL-3’-Cy5 to the EF-G labeled at F410C position with Cy3 dye, indicating proper interaction between the two molecules. Stars showed the positions of dye labeling. The particle counts for the FRET efficiency histograms can be found in **Table S1**.

**Figure S3.**
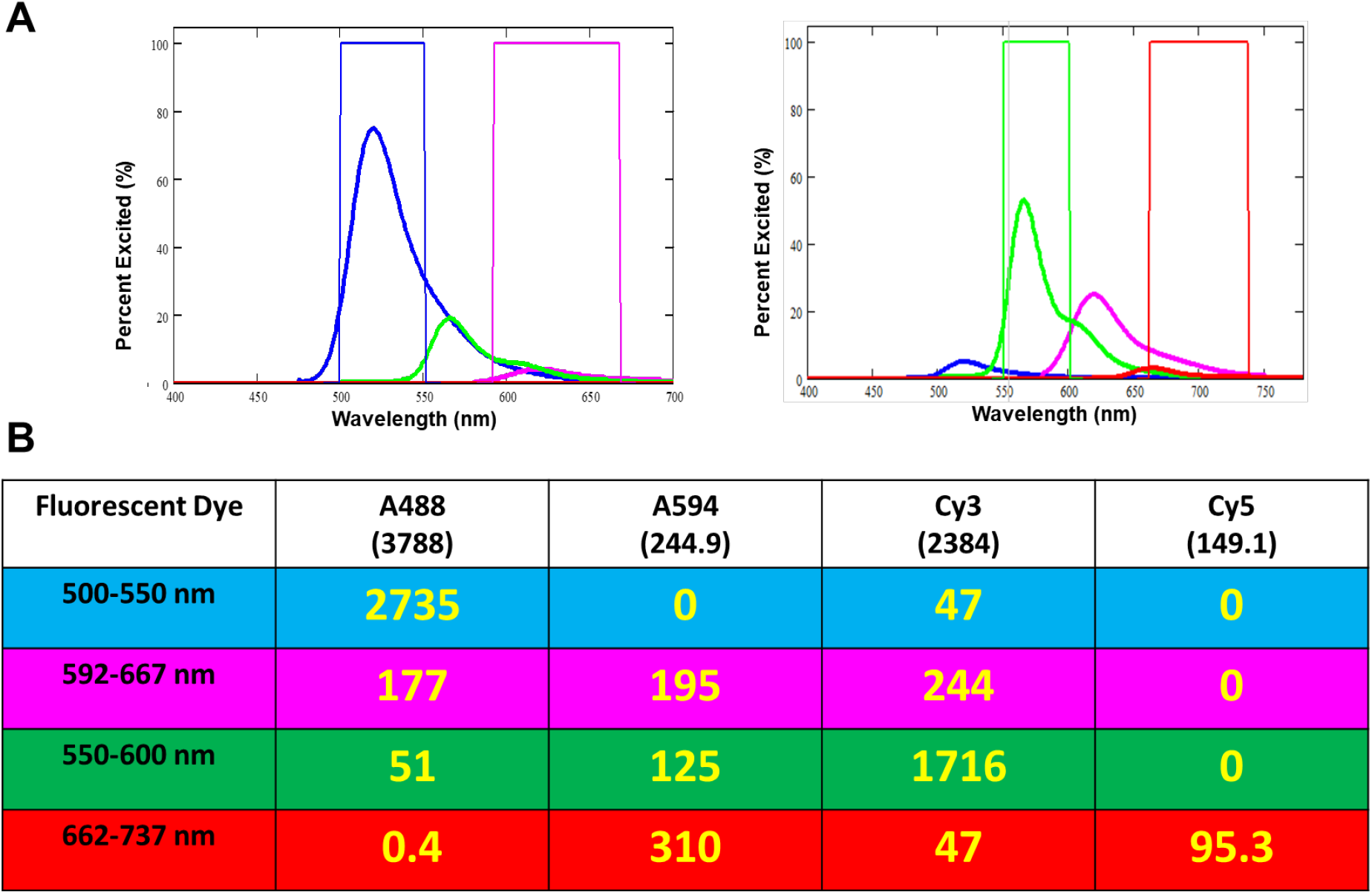
Crosstalk correction procedure for dual channel smFRET experiments. **(A)** Fluorescence emission spectra of all fluorophores under 488 nm (left) or 532 nm (right) laser excitation. The intensities are corrected by laser absorption and quantum yields. Colored rectangles show filters used for the Alex488 (blue)/Alexa594 (magenta) and Cy3 (green)/Cy5 pairs (red), respectively. **(B)** Relative fluorescence intensities for dyes within the filters as shown in **(A)**. The intensities are scaled by excitation efficiencies under the specific laser line, after considering the absorption and emission yields (https://www.thermofisher.com/order/fluorescence-spectraviewer#!/).

Crosstalk between laser wavelengths and filter sets was observed during multi-channel smFRET experiments. There were undesired direct excitations and spectrum crosstalks. For example, in **Figure S3A**, Cy3 direct excitation (green trace) by 488 nm laser spilled over into 488/594 FRET filter windows (blue and magenta rectangles). Similarly, Alexa 488 and Alexa 594 dyes were direct excited by 532 nm laser and bled into Cy3/Cy5 FRET filter windows (green and red rectangles). The composition of signal in each channel under both laser excitations were tabulated in **Figure S3B** and deconvoluted in FRET calculation. The values in **Figure S3B** are the integration area beneath the fluorescence emission spectra defined by the filter boundaries. The values in the parenthesis are the total area for each fluorescence curve.

#### Deconvolution for Alexa488/594 FRET pair under 488 nm laser excitation

The second and third rows in the above table indicate that the sum of fluorescence intensities from both the donor and acceptor channels is 2735 + 47 + 177 + 195 + 244, totaling 3398. According to FRET theory, even though the intensities of the donor and acceptor can change due to energy transfer, their combined fluorescence should always match the initial intensity of the donor alone. To illustrate, a FRET efficiency of 50% would result in 50% * 2735 in the donor channel and the same amount in the acceptor channel. Similarly, a 20% FRET efficiency would yield 80% * 2735 in the donor channel and 20% * 2735 in the acceptor channel. In each scenario, the combined fluorescence from the two channels remains unchanged, regardless of the FRET efficiency. By considering this combined fluorescence, the variable FRET signals across donor and acceptor channels are eliminated from the formula. For clarity, ’SUM’ denotes the experimental total fluorescence intensities of both channels, while ’D’ and ’A’ represent the experimental fluorescence intensity in the donor and acceptor channels, respectively. Consequently, I_A488_ = D-SUM*47/3398 = D-SUM*1.38%. Here, deconvolute with “SUM*47/3398” is more accurate than use “D*47/2735” because the area 2735 represent the theoretical total energy that will be partitioned between the donor and acceptor, while D is the experimental value after FRET occurred. Given that the experimental D value doesn’t scale linearly with the theoretical 2735 value due to energy transferred away to the acceptor, the contamination from Cy3 isn’t proportionate to the 47/2735 factor. However, irrespective of FRET, the combined energy from both the donor and acceptor channels should align proportionally with the theoretical sum values, as we elaborated above. As a result, the scaling factors are determined using the SUM as the denominator. Similarly, I_A594_ = A-SUM*(177+195+244)/3398=A-SUM*18%. So FRET is calculated by (I_A594_)/(I_A594_ + I_A488_) = **(82%*A-18%D)/(SUM*81.6%)**

#### Deconvolution for *Cy3*/*Cy5* FRET pair under *532* nm laser excitation

*Using similar formula, I_Cy3_ =* D-SUM*(51+125)/2280.7 = D-SUM*7.7%. Similarly, *I_Cy5_ =* A-SUM*(0.4+310+47+95.3)/2280.7=A-SUM*20%. So FRET is calculated by (I_Cy5_)/(I_Cy5_ + I_Cy3_) = **(80%*A-20%D)/(SUM*72.3%)**

**Figure S4.**
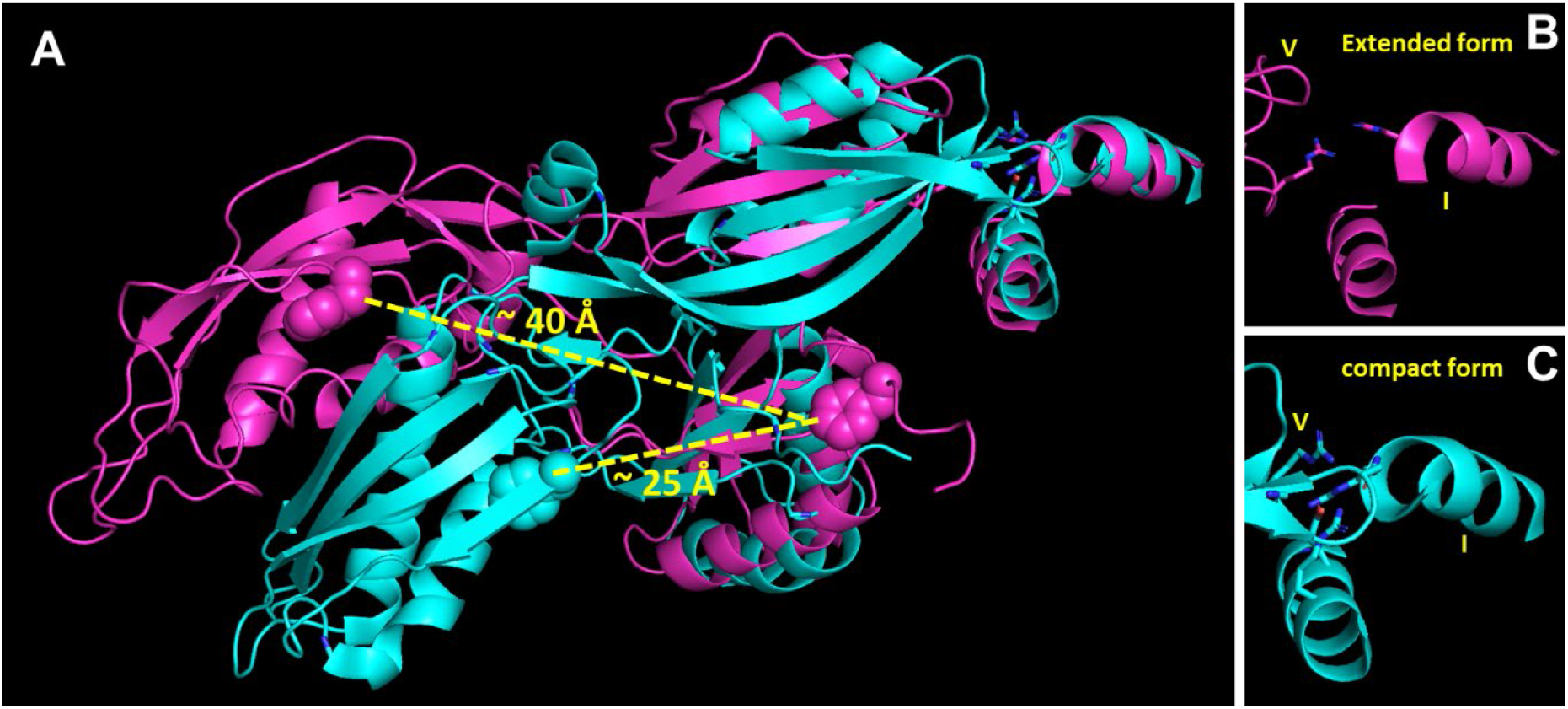
Examples of extended and compact EF-G conformations. **(A)** Overlay of free EF-G (1fnm, cyan) and ribosome bound EF-G (7pjv, magenta). **(B-C)** The extension of the domain VI is accompanied by an increased distance between domain I and V. The residues 410 and 533 are depicted in spheres form.

**Figure S5.**
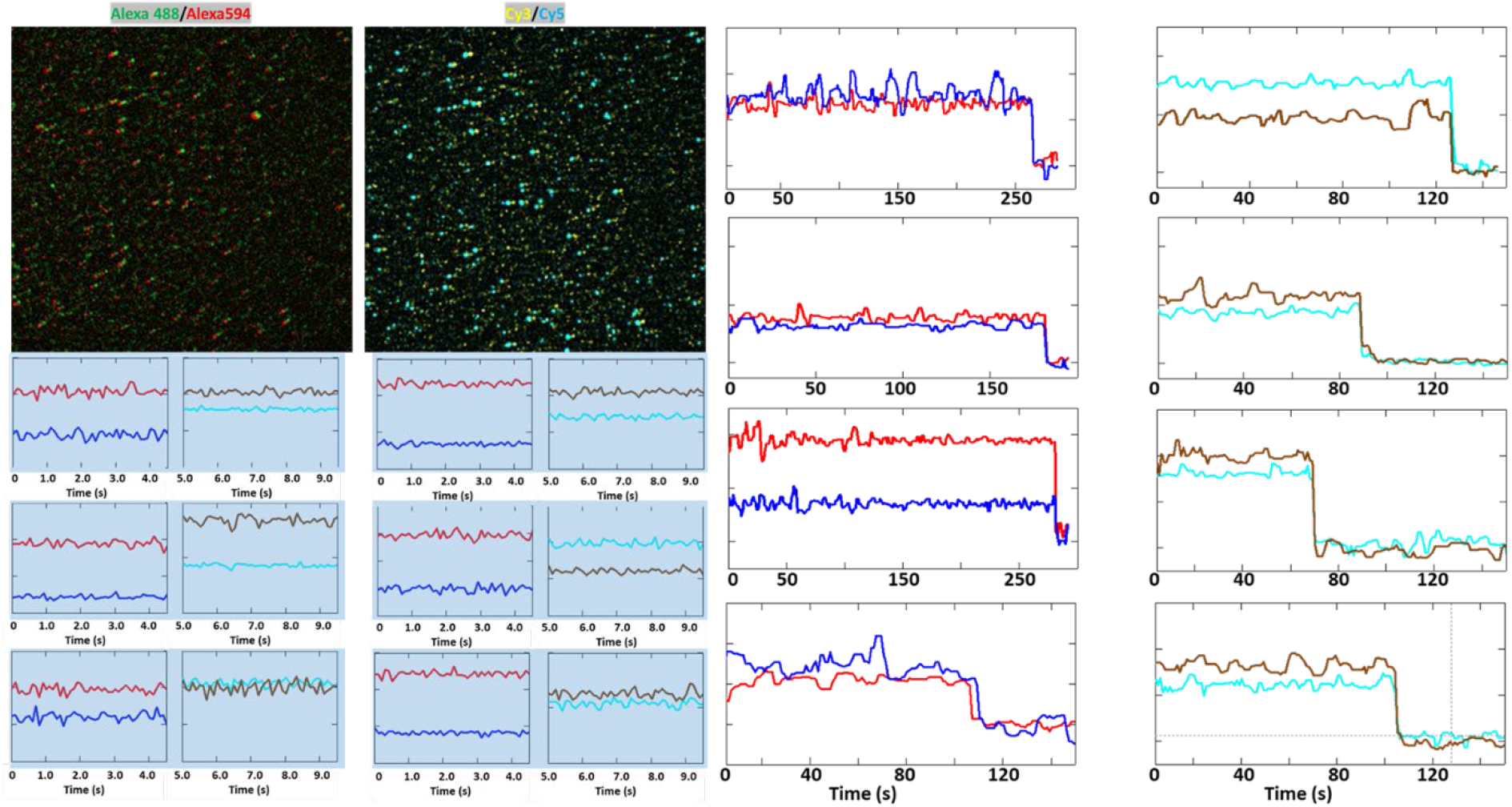
Representative image and fitted fluorescence intensity traces from Figure 5J histogram (total particles 2328). Red/Blue: Alexa 488/Alexa 594; Brown/Cyan: Cy3/Cy5.

**Figure S5** displays some representative smFRET images and some fitted traces. Single step bleaching traces were collected at prolonged time-lapse, and 10x higher laser power (400 mW) was exerted to bleach the Cy3/Cy5 FRET pair.

The left camera image shows an overlay emission of Alexa488/Alex a594 that are labeled on EF-G, while the right camera image is an overlay emission of Cy3/Cy5 that are labeled on tRNA/L27 (time-lapse traces are shorted for display clarity). They are the same ribosome complexes at the matching coordinates. These images are stacked into one movie, and the fluorescence intensities were fitted in one trace of 100 time points, in which the first 50 points reflect signals from the Alexa488/Alexa594 channels, and the subsequent 50 points depict signals from the Cy3/Cy5 channels. The single molecule particles are selected by program with consistent parameters. The selection criteria mirror those previously described (1). Specifically, intensity values are confined between 100 and 2000, and the signal-to-noise ratio (S/N) must exceed 4. Traces that meet these stipulations across all four channels are chosen for further analysis. For instance, in **Figure 5J**, 6626 particles were chosen from the EF-G FRET images based on the first 5 seconds of the fitted traces. Of these, 2328 particles demonstrated signals surpassing the threshold from the Cy3/Cy5 FRET images during the subsequent 5 seconds. Consequently, the histogram was compiled from this final collection of 2328 particles. The particle counts for the FRET efficiency histograms can be found in **Table S1**.

**Figure S6.**
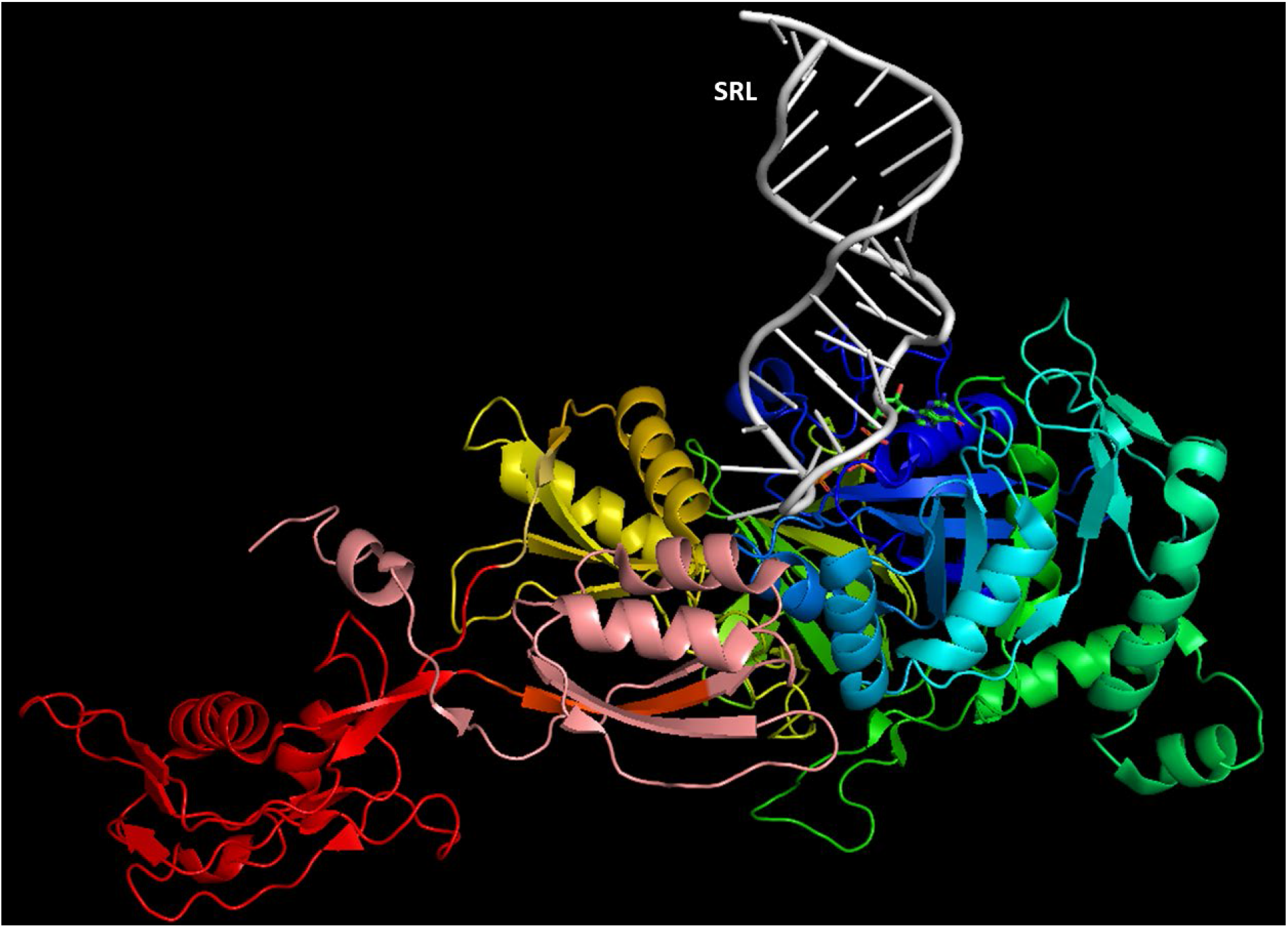
Interaction of SR loop with the EF-G (based on structure 7pjv).

**Figure S7.**
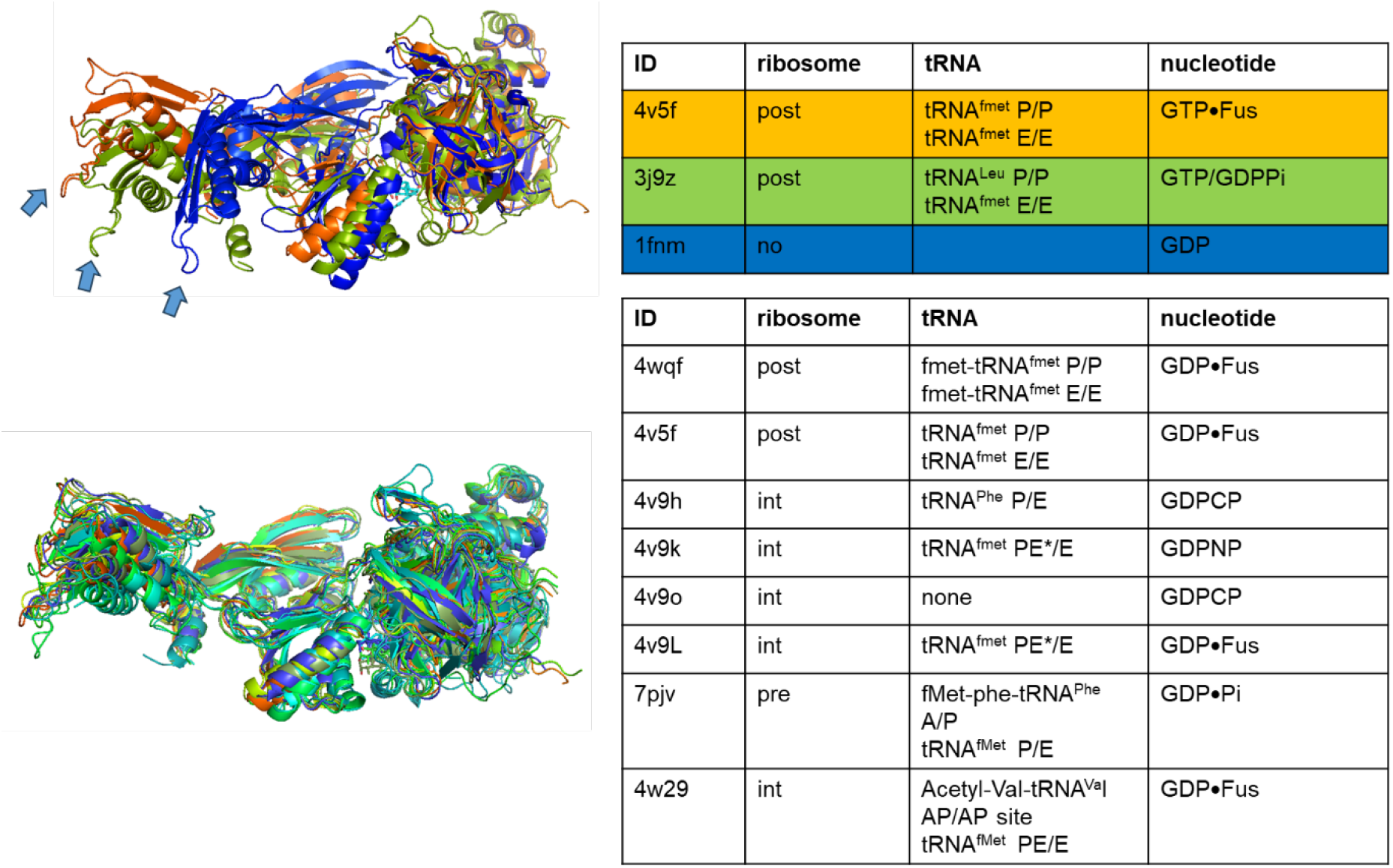
Structure variations of EF-G in bound and free forms (1fnm). The top panel overlaid three representative structures of EF-G, with increasing distances between the domain IV and domain III, in the order of 1fnm, 3j9z, and 4v5f. The bottom panel showcases some of the representative bound EF-G complexes with similar extended conformation, irrespective of tRNA and nucleotide configurations.

**Figure S8.**
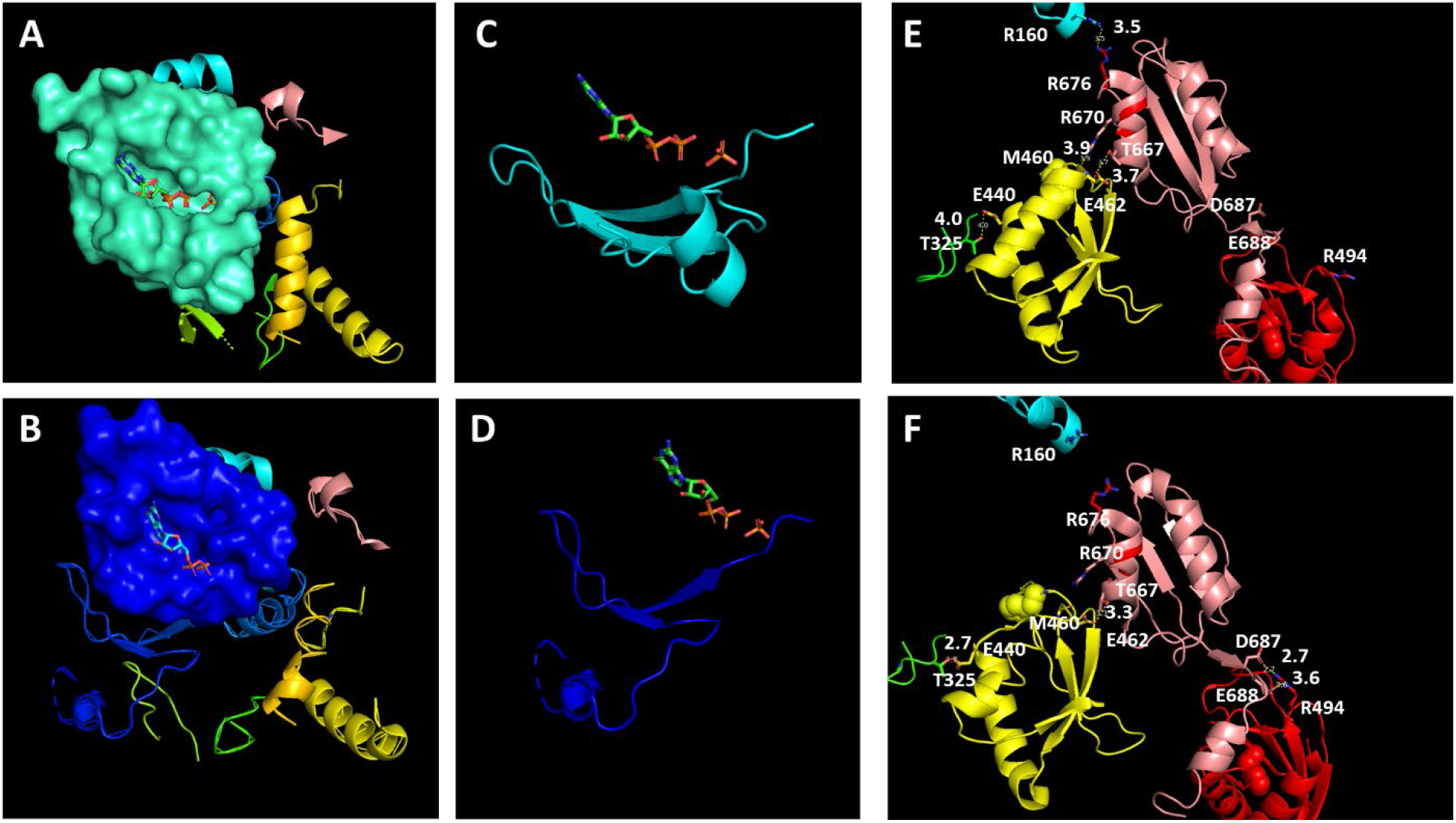
**(A)** The closed GTP binding pocket before Pi releasing but after GTP hydrolysis (7pjv). **(B)** The opened GTP binding pocket after Pi releasing (7pjy). **(C)** The effect loop and GDP•Pi in the closed form. **(D)** The effect loop and GDP•Pi in the opened form. The Pi is overlay from (C). **(E-F)** Interdomain interactions of domain III (yellow), IV (red) and V (salmon) to show the molecular rearrangement during the opening process. Some of the residues are shown to break or form interdomain H-bondings.

**Figure S9.**
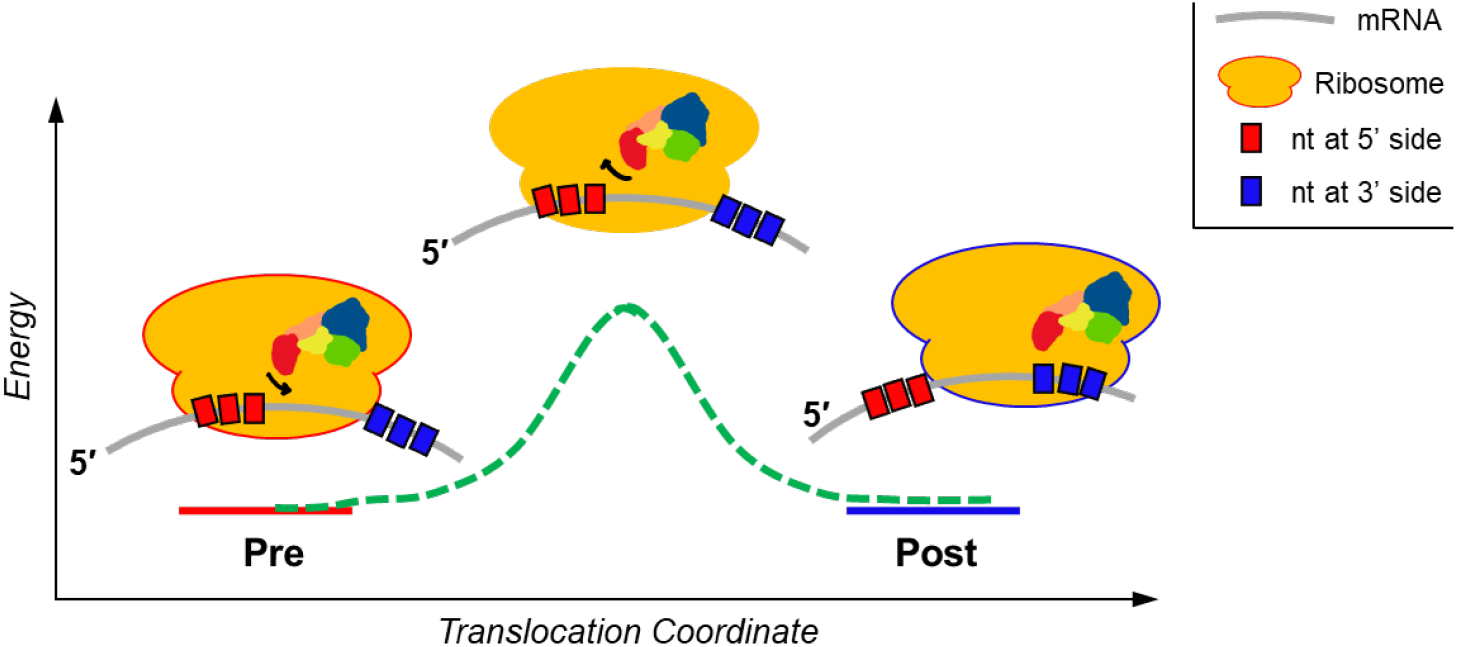
A illustration of the possible role of the highly compact EF-G during translocation.

**Figure S10.**
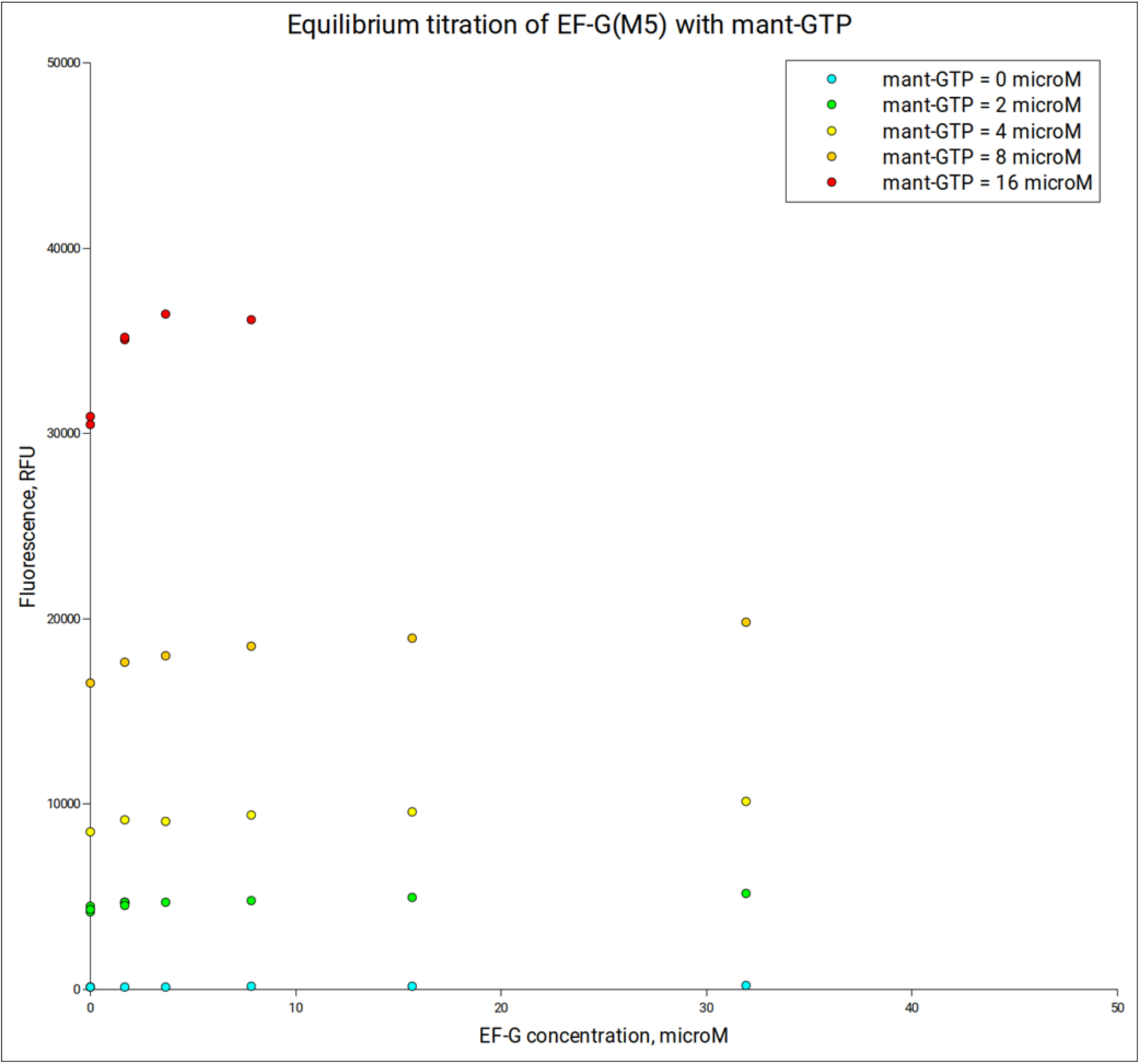
Equilibrium titration of EF-G (M5) with mant-GTP. Experiments were performed at room temperature (20°C) with samples prepared in Dilution buffer (50 mM Tris-HCl, pH 7.5, 70 mM NH4Cl, 30 mM KCl, 10 mM MgCl2, 0.5 mM EDTA, 6 mM 2-mercaptoethanol). Fluorescence was measured using a DeNovix QFX fluorometer set with excitation wavelengths in 361-389 nm range and fluorescence detection in 435-485 nm range. Each plotted value represents an average of three measurements.

**Figure S11.**
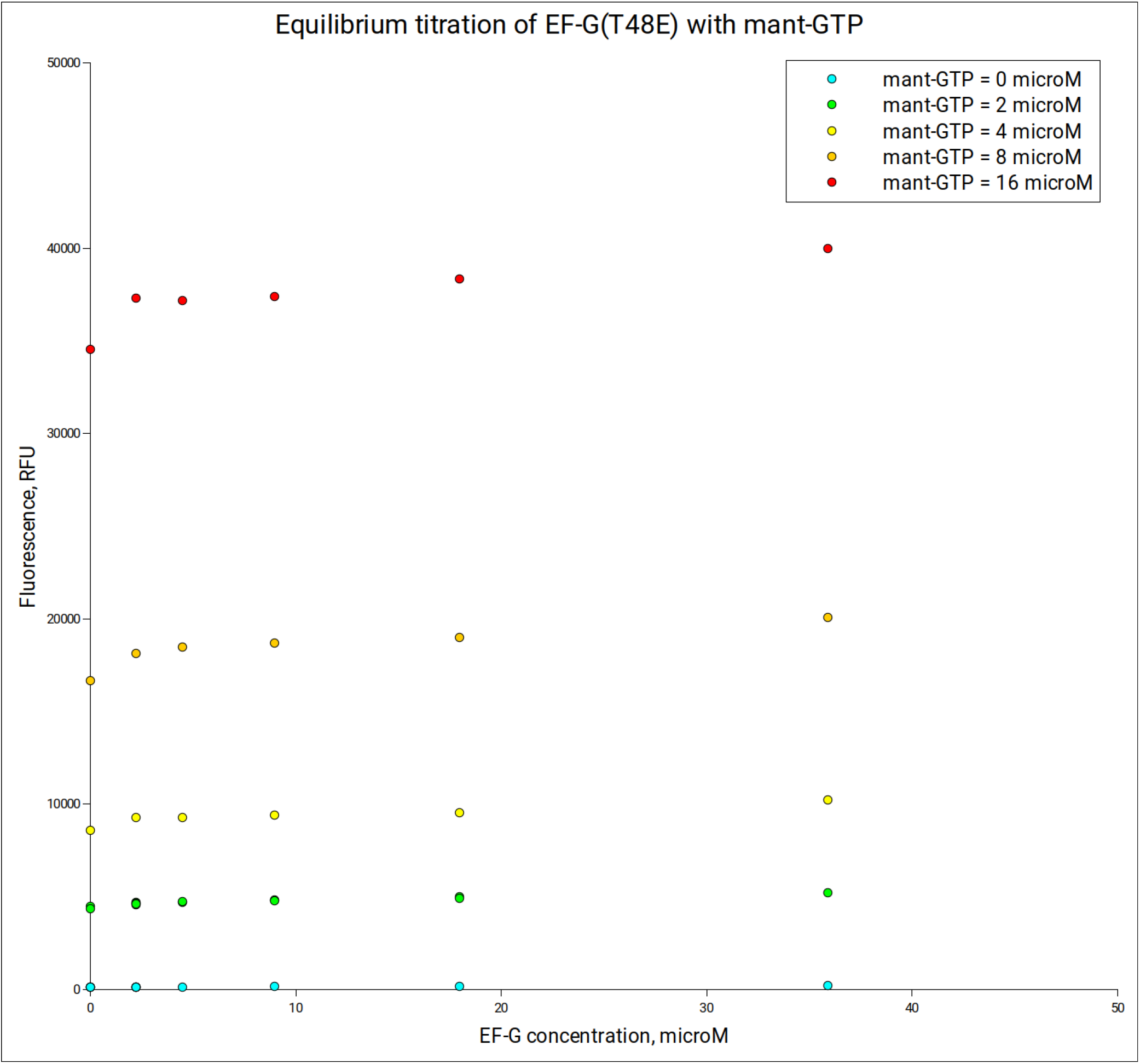
Equilibrium titration of EF-G (T48E) with mant-GTP. Experiments were performed at room temperature (20°C) with samples prepared in Dilution buffer (50 mM Tris-HCl, pH 7.5, 70 mM NH4Cl, 30 mM KCl, 10 mM MgCl2, 0.5 mM EDTA, 6 mM 2-mercaptoethanol). Fluorescence was measured using a DeNovix QFX fluorometer set with excitation wavelengths in 361-389 nm range and fluorescence detection in 435-485 nm range. Each plotted value represents an average of three measurements.

**Figure S12.**
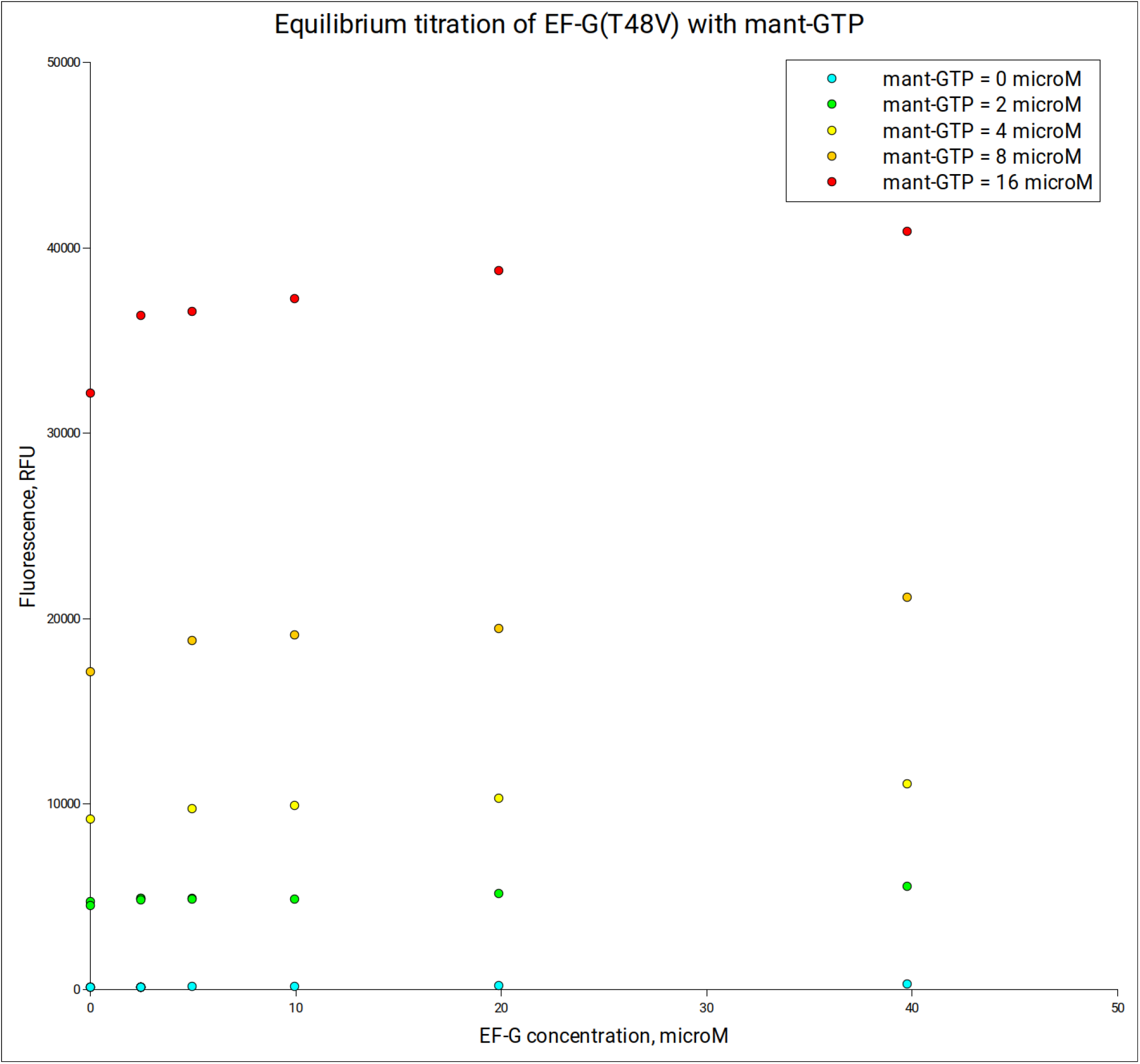
Equilibrium titration of EF-G (T48V) with mant-GTP. Experiments were performed at room temperature (20°C) with samples prepared in Dilution buffer (50 mM Tris-HCl, pH 7.5, 70 mM NH_4_Cl, 30 mM KCl, 10 mM MgCl_2_, 0.5 mM EDTA, 6 mM 2-mercaptoethanol). Fluorescence was measured using a DeNovix QFX fluorometer set with excitation wavelengths in 361-389 nm range and fluorescence detection in 435-485 nm range. Each plotted value represents an average of three measurements.

**Figure S13.**
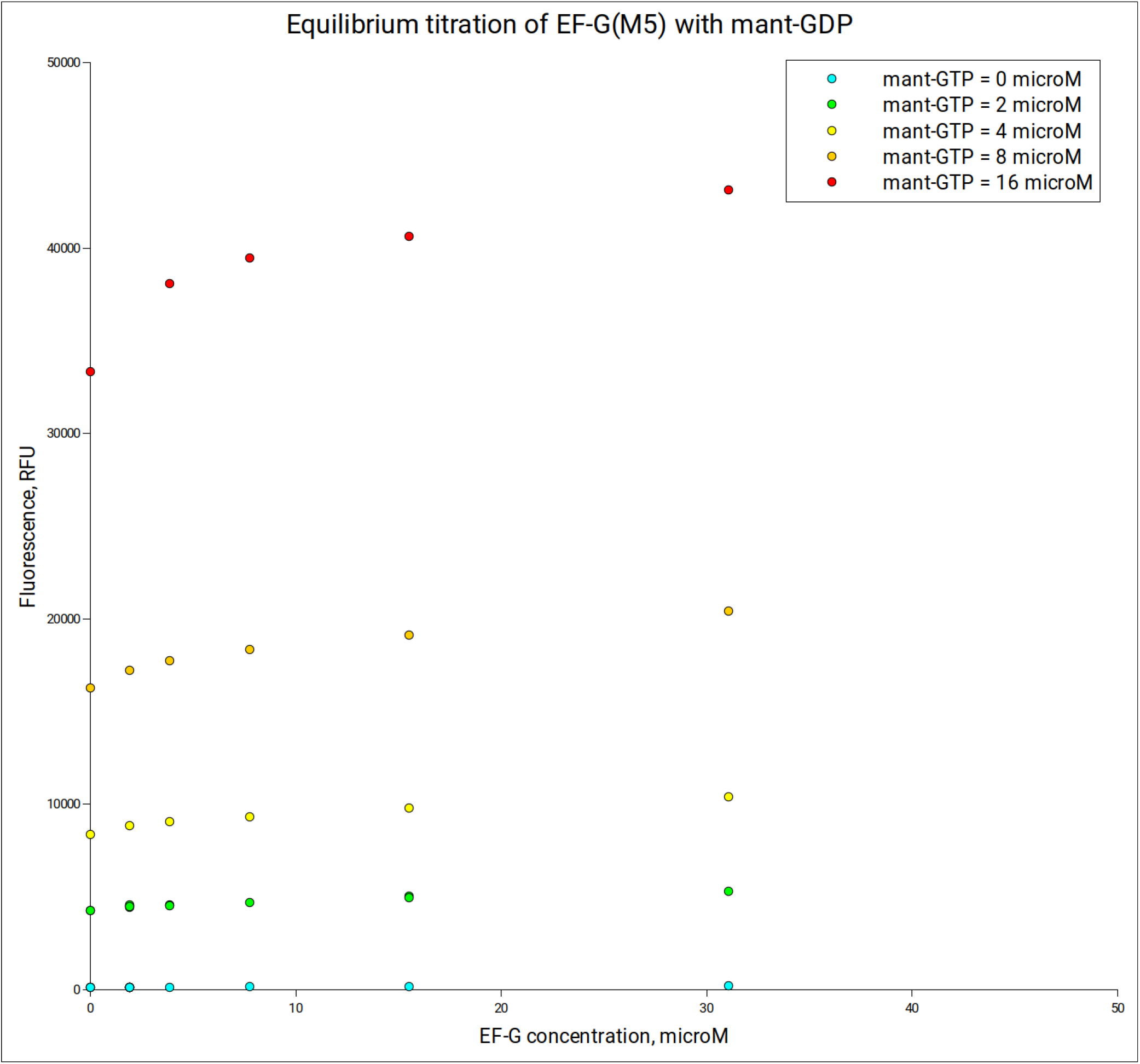
Equilibrium titration of EF-G (M5) with mant-GDP. Experiments were performed at room temperature (20°C) with samples prepared in Dilution buffer (50 mM Tris-HCl, pH 7.5, 70 mM NH_4_Cl, 30 mM KCl, 10 mM MgCl_2_, 0.5 mM EDTA, 6 mM 2-mercaptoethanol). Fluorescence was measured using a DeNovix QFX fluorometer set with excitation wavelengths in 361-389 nm range and fluorescence detection in 435-485 nm range. Each plotted value represents an average of three measurements.

**Figure S14.**
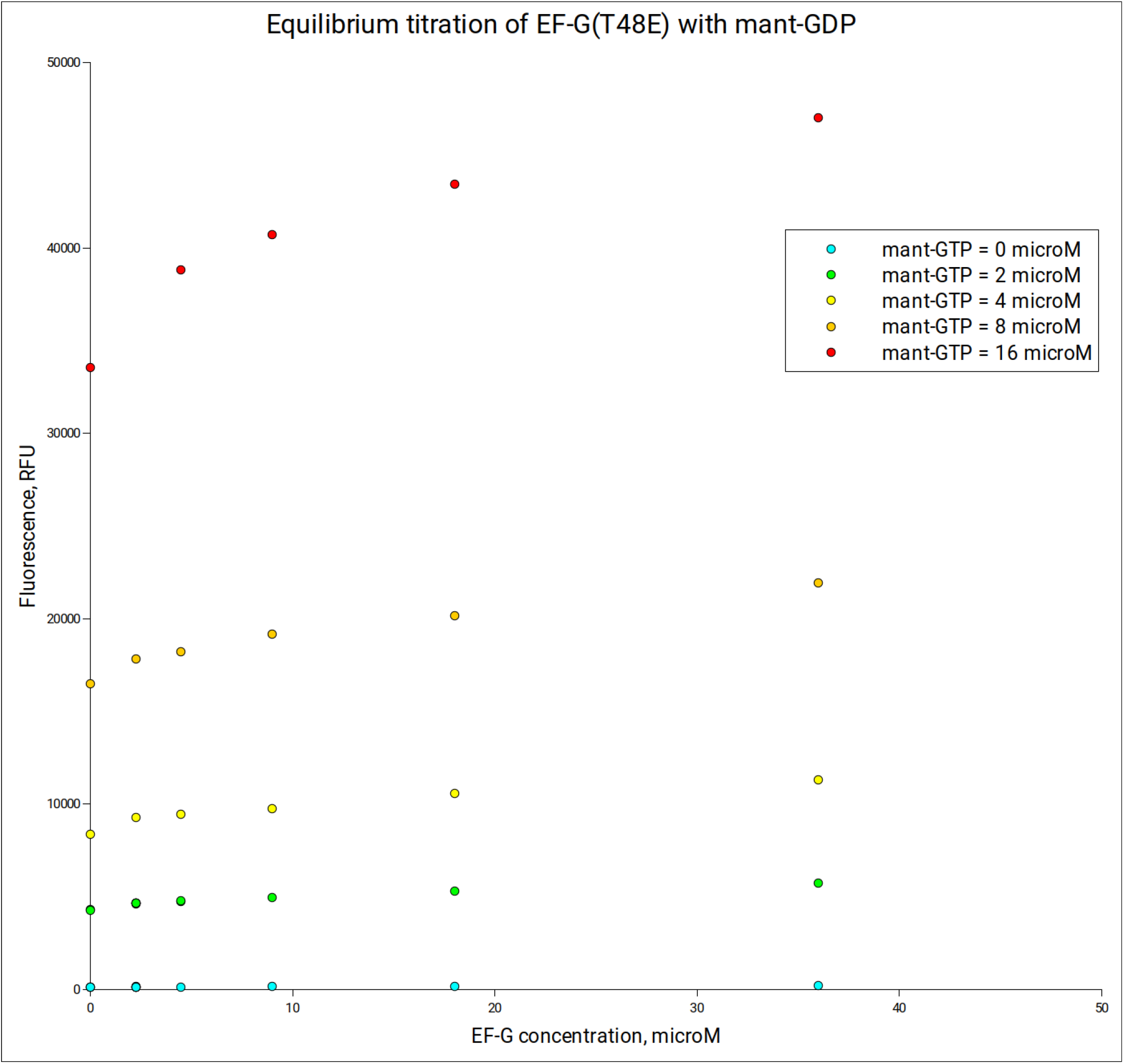
Equilibrium titration of EF-G (T48E) with mant-GDP. Experiments were performed at room temperature (20°C) with samples prepared in Dilution buffer (50 mM Tris-HCl, pH 7.5, 70 mM NH_4_Cl, 30 mM KCl, 10 mM MgCl_2_, 0.5 mM EDTA, 6 mM 2-mercaptoethanol). Fluorescence was measured using a DeNovix QFX fluorometer set with excitation wavelengths in 361-389 nm range and fluorescence detection in 435-485 nm range. Each plotted value represents an average of three measurements.

**Figure S15.**
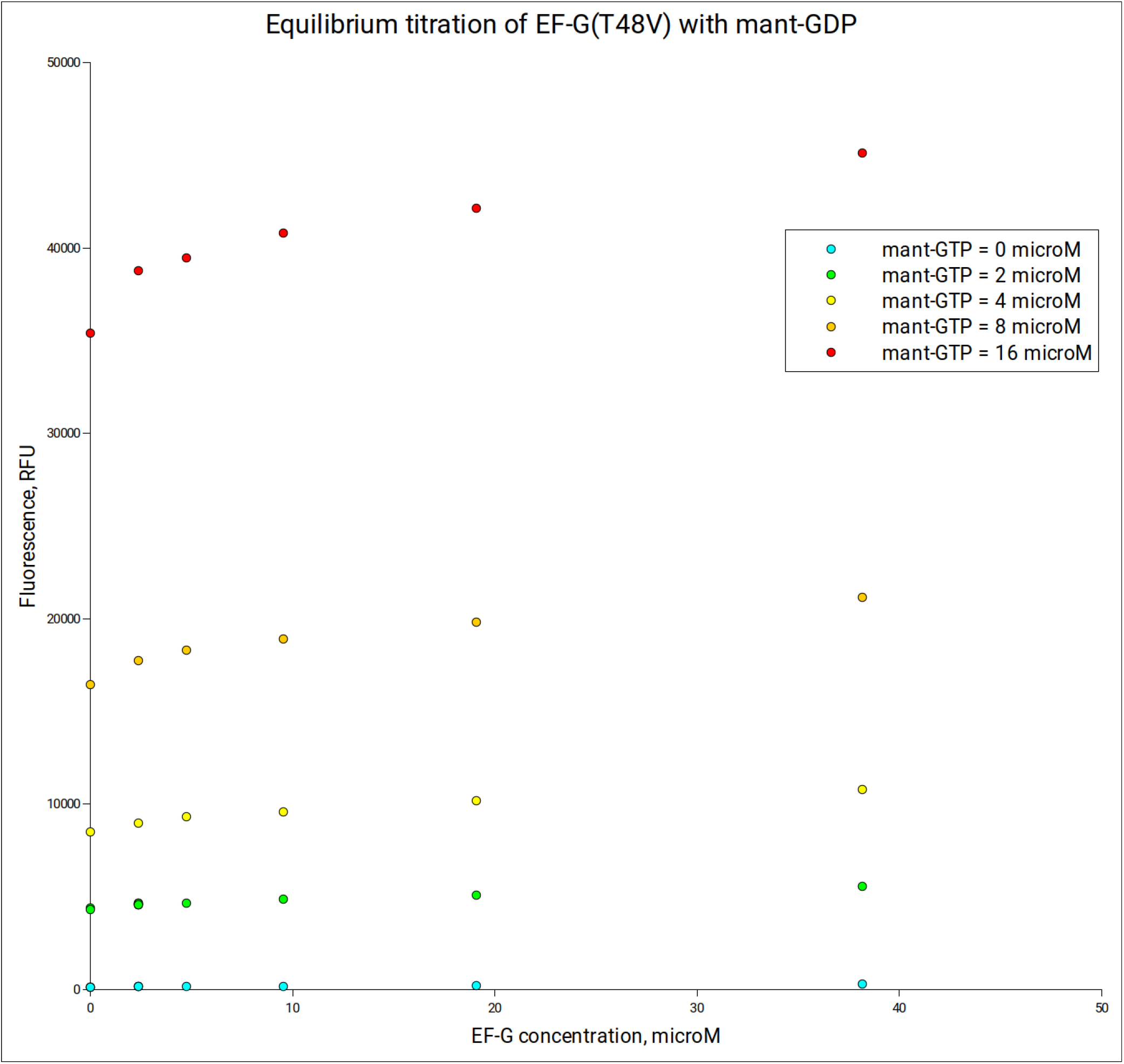
Equilibrium titration of EF-G (T48V) with mant-GDP. Experiments were performed at room temperature (20°C) with samples prepared in Dilution buffer (50 mM Tris-HCl, pH 7.5, 70 mM NH_4_Cl, 30 mM KCl, 10 mM MgCl_2_, 0.5 mM EDTA, 6 mM 2-mercaptoethanol). Fluorescence was measured using a DeNovix QFX fluorometer set with excitation wavelengths in 361-389 nm range and fluorescence detection in 435-485 nm range. Each plotted value represents an average of three measurements.

## REFERENCES

1. T. M. Schmeing, V. Ramakrishnan, What recent ribosome structures have revealed about the mechanism of translation. Nature 461, 1234–1242 (2009).

2. S. R. Connell et al., Structural basis for interaction of the ribosome with the switch regions of GTP-bound elongation factors. Mol Cell 25, 751–764 (2007).

3. J. S. Gonzalez-Garcia, A model for ribosome translocation based on the alternated displacement of its subunits. Eur Biophys J 52, 175–187 (2023).

4. B. S. Schuwirth et al., Structures of the bacterial ribosome at 3.5 A resolution. Science 310, 827–834 (2005).

5. J. Frank, R. K. Agrawal, A ratchet-like inter-subunit reorganization of the ribosome during translocation. Nature 406, 318–322 (2000).

6. M. V. Rodnina, A. Savelsbergh, V. I. Katunin, W. Wintermeyer, Hydrolysis of GTP by elongation factor G drives tRNA movement on the ribosome. Nature 385, 37–41 (1997).

7. J. Zhou, L. Lancaster, J. P. Donohue, H. F. Noller, Spontaneous ribosomal translocation of mRNA and tRNAs into a chimeric hybrid state. Proc Natl Acad Sci U S A 116, 7813–7818 (2019).

8. P. V. Cornish, D. N. Ermolenko, H. F. Noller, T. Ha, Spontaneous intersubunit rotation in single ribosomes. Mol Cell 30, 578–588 (2008).

9. J. Lin, M. G. Gagnon, D. Bulkley, T. A. Steitz, Conformational changes of elongation factor G on the ribosome during tRNA translocation. Cell 160, 219–227 (2015).

10. H. Stark, M. V. Rodnina, H. J. Wieden, M. van Heel, W. Wintermeyer, Large-scale movement of elongation factor G and extensive conformational change of the ribosome during translocation. Cell 100, 301–309 (2000).

11. W. Li et al., Activation of GTP hydrolysis in mRNA-tRNA translocation by elongation factor G. Sci Adv 1 (2015).

12. V. Petrychenko et al., Structural mechanism of GTPase-powered ribosome-tRNA movement. Nat Commun 12, 5933 (2021).

13. C. E. Carbone et al., Time-resolved cryo-EM visualizes ribosomal translocation with EF-G and GTP. Nat Commun 12, 7236 (2021).

14. C. E. Cunha et al., Dual use of GTP hydrolysis by elongation factor G on the ribosome. Translation (Austin) 1, e24315 (2013).

15. W. Holtkamp et al., GTP hydrolysis by EF-G synchronizes tRNA movement on small and large ribosomal subunits. EMBO J 33, 1073–1085 (2014).

16. J. Chen, A. Petrov, A. Tsai, S. E. O’Leary, J. D. Puglisi, Coordinated conformational and compositional dynamics drive ribosome translocation. Nat Struct Mol Biol 20, 718–727 (2013).

17. L. Yao, Y. Li, T. W. Tsai, S. Xu, Y. Wang, Noninvasive measurement of the mechanical force generated by motor protein EF-G during ribosome translocation. Angewandte Chemie 52, 14041–14044 (2013).

18. H. Yin, M. Gavriliuc, R. Lin, S. Xu, Y. Wang, Modulation and Visualization of EF-G Power Stroke During Ribosomal Translocation. Chembiochem 20, 2927–2935 (2019).

19. G. Rexroad, J. P. Donohue, L. Lancaster, H. F. Noller, The role of GTP hydrolysis by EF-G in ribosomal translocation. Proc Natl Acad Sci U S A 119, e2212502119 (2022).

20. J. Frank, R. L. Gonzalez, Jr., Structure and dynamics of a processive Brownian motor: the translating ribosome. Annu Rev Biochem 79, 381–412 (2010).

21. E. Salsi, E. Farah, D. N. Ermolenko, EF-G Activation by Phosphate Analogs. J Mol Biol 428, 2248–2258 (2016).

22. E. J. Rundlet et al., Structural basis of early translocation events on the ribosome. Nature 595, 741–745 (2021).

23. R. Liu, C. G. Proud, Eukaryotic elongation factor 2 kinase as a drug target in cancer, and in cardiovascular and neurodegenerative diseases. Acta Pharmacol Sin 37, 285–294 (2016).

24. A. G. Ryazanov, E. K. Davydova, Mechanism of elongation factor 2 (EF-2) inactivation upon phosphorylation. Phosphorylated EF-2 is unable to catalyze translocation. FEBS Lett 251, 187–190 (1989).

25. A. G. Ryazanov, E. A. Shestakova, P. G. Natapov, Phosphorylation of elongation factor 2 by EF-2 kinase affects rate of translation. Nature 334, 170–173 (1988).

26. S. Majumdar et al., Insights into translocation mechanism and ribosome evolution from cryo-EM structures of translocation intermediates of Giardia intestinalis. Nucleic Acids Res 51, 3436–3451 (2023).

27. K. S. Wilson, H. F. Noller, Mapping the position of translational elongation factor EF-G in the ribosome by directed hydroxyl radical probing. Cell 92, 131–139 (1998).

28. M. E. Altuntop, C. T. Ly, Y. Wang, Single-molecule study of ribosome hierarchic dynamics at the peptidyl transferase center. Biophysical journal 99, 3002–3009 (2010).

29. B. A. Maguire, A. D. Beniaminov, H. Ramu, A. S. Mankin, R. A. Zimmermann, A protein component at the heart of an RNA machine: the importance of protein l27 for the function of the bacterial ribosome. Mol Cell 20, 427–435 (2005).

30. D. Pan, H. Qin, B. S. Cooperman, Synthesis and functional activity of tRNAs labeled with fluorescent hydrazides in the D-loop. RNA 15, 346–354 (2009).

31. M. R. Wasserman, J. L. Alejo, R. B. Altman, S. C. Blanchard, Multiperspective smFRET reveals rate-determining late intermediates of ribosomal translocation. Nat Struct Mol Biol 23, 333–341 (2016).

32. M. K. Yang et al., Global phosphoproteomic analysis reveals diverse functions of serine/threonine/tyrosine phosphorylation in the model cyanobacterium Synechococcus sp. strain PCC 7002. J Proteome Res 12, 1909–1923 (2013).

33. B. Wilden, A. Savelsbergh, M. V. Rodnina, W. Wintermeyer, Role and timing of GTP binding and hydrolysis during EF-G-dependent tRNA translocation on the ribosome. Proc Natl Acad Sci U S A 103, 13670–13675 (2006).

34. J. Verhelst et al., Functional Comparison of Mx1 from Two Different Mouse Species Reveals the Involvement of Loop L4 in the Antiviral Activity against Influenza A Viruses. J Virol 89, 10879–10890 (2015).

35. Y. Endo, P. W. Huber, I. G. Wool, The ribonuclease activity of the cytotoxin alpha-sarcin. The characteristics of the enzymatic activity of alpha-sarcin with ribosomes and ribonucleic acids as substrates. J Biol Chem 258, 2662–2667 (1983).

36. T. P. Hausner, J. Atmadja, K. H. Nierhaus, Evidence that the G2661 region of 23S rRNA is located at the ribosomal binding sites of both elongation factors. Biochimie 69, 911–923 (1987).

37. X. Shi, P. K. Khade, K. Y. Sanbonmatsu, S. Joseph, Functional role of the sarcin-ricin loop of the 23S rRNA in the elongation cycle of protein synthesis. J Mol Biol 419, 125–138 (2012).

38. B. Zhang et al., Progress in the Development of Eukaryotic Elongation Factor 2 Kinase (eEF2K) Natural Product and Synthetic Small Molecule Inhibitors for Cancer Chemotherapy. Int J Mol Sci 22 (2021).

39. N. P. Kasica et al., Antagonists targeting eEF2 kinase rescue multiple aspects of pathophysiology in Alzheimer’s disease model mice. J Neurochem 160, 524–539 (2022).

40. C. Heise et al., Elongation factor-2 phosphorylation in dendrites and the regulation of dendritic mRNA translation in neurons. Front Cell Neurosci 8, 35 (2014).

41. E. Salsi, E. Farah, Z. Netter, J. Dann, D. N. Ermolenko, Movement of elongation factor G between compact and extended conformations. J Mol Biol 427, 454–467 (2015).

## Reference

1. M. E. Altuntop, C. T. Ly, Y. Wang, Single-molecule study of ribosome hierarchic dynamics at the peptidyl transferase center. Biophysical journal 99, 3002–3009 (2010).

2. T. W. Tsai, H. Yang, H. Yin, S. Xu, Y. Wang, High-Efficiency "-1" and "-2" Ribosomal Frameshiftings Revealed by Force Spectroscopy. ACS chemical biology 12, 1629–1635 (2017).

3. J. S. Dubnoff, U. Maitra, Isolation and properties of polypeptide chain initiation factor FII from Escherichia coli: evidence for a dual function. Proc Natl Acad Sci U S A 68, 318–323 (1971).

4. A. L. Haenni, F. Chapeville, The behaviour of acetylphenylalanyl soluble ribonucleic acid in polyphenylalanine synthesis. Biochim Biophys Acta 114, 135–148 (1966).

5. B. Wilden, A. Savelsbergh, M. V. Rodnina, W. Wintermeyer, Role and timing of GTP binding and hydrolysis during EF-G-dependent tRNA translocation on the ribosome. Proc Natl Acad Sci U S A 103, 13670–13675 (2006).

6. J. Verhelst et al., Functional Comparison of Mx1 from Two Different Mouse Species Reveals the Involvement of Loop L4 in the Antiviral Activity against Influenza A Viruses. J Virol 89, 10879–10890 (2015).

7. A. Pulk, J. H. Cate, Control of ribosomal subunit rotation by elongation factor G. Science 340, 1235970 (2013).

